# Trifunctional fatty acid derivatives demonstrate the impact of diazirine placement

**DOI:** 10.1101/2024.05.15.594383

**Authors:** Scotland E Farley, Ryu Hashimoto, Judah Evangelista, Frank Stein, Per Haberkant, Kazuya Kikuchi, Fikadu G. Tafesse, Carsten Schultz

**Affiliations:** Oregon Health & Science University, Department of Chemical Physiology and Biochemistry, Portland, OR, 97239, USA; Oregon Health & Science University, Department of Molecular Microbiology and Immunology, Portland, OR, 97239, USA; Osaka University, Department of Applied Chemistry, Graduate School of Engineering, 2-1 Yamadaoka, Suita, Osaka, 565-0871, Japan; European Molecular Biology Laboratory, Proteomics Core Facility, 69117, Heidelberg, Germany

## Abstract

Functionalized lipid probes are a critical new tool to interrogate the function of individual lipid species, but the structural parameters that constrain their utility have not been thoroughly described. Here, we synthesize three palmitic acid derivatives with a diazirine at different positions on the acyl chain and examine their metabolism, subcellular localization, and protein interactions. We demonstrate that while they produce very similar metabolites and subcellular distributions, probes with the diazirine at either end pulldown distinct subsets of proteins after photo-crosslinking. This highlights the importance of thoughtful diazirine placement when developing probes based on biological molecules.

## RESULTS AND DISCUSSION

Historically, it has been extremely difficult to elucidate the subcellular role of individual lipids, due to the scarcity of tools to directly observe them. Lipids are difficult to genetically manipulate, as the knockout of specific biosynthetic enzymes often leads to the ablation of a class of lipids rather than an individual species, and even that is often accompanied by unpredictable compensatory effects elsewhere in the lipidome^1^. Furthermore, lipids cannot be genetically tagged in the way that proteins can, and there are very few antibodies or sensors for single lipid species^2^.

To overcome these difficulties, chemical biology strategies have been adapted to the study of lipids over the last decade, including bioorthogonal labeling^3-4^, photo-crosslinking^5^, and photocaging^6^. Lipid-based chemical probes require several features to be biologically useful: they should be taggable (for visualization or affinity purification), photo-crosslinkable (to identify binding partners), cell-permeant, transiently inert to cellular metabolism (via a caging group), and, of course, faithful to the biological properties of the lipids they represent. While caged lipid derivatives^6^ and probes with a diazirine and an alkyne^5^ were published earlier, a strategy to achieve probes with all three features was described for a trifunctional sphingosine in 2015, and an additional fatty acid, and diacylglycerol in 2017^8^. These latter studies of trifunctional lipid probes demonstrated that the trifunctional lipid derivatives could recapitulate key aspects of the parent lipids’ biology, and that the two photoactive groups were chromatically orthogonal: blue light (400 nm) that could photocleave the coumarin cage would not activate the UV-reactive (350 nm) diazirine^8^. Since then, the strategy has been applied to a variety of lipids, including phosphatidylinositol^9^, PI(3,4)P_2_, PI(3,4,5)P_3_^10^, and sphinganine^11^. In all probes, the diazirine was placed at the C10 position of a fatty acid (8-3 fatty acid, **1**, Scheme 1B) or at a similar distance to the headgroup in sphingolipids. This raised questions about what lipid binding proteins could be captured by the crosslinking reaction. We therefore wanted to investigate how the position of the diazirine on the lipid backbone affects the metabolism, the cellular location and protein interactions of the resulting lipid. Diazirine placement could impact lipid probe function in several ways: 1) by altering its recognition by lipid metabolic enzymes, resulting in a different collection of probe-derived lipid metabolites; 2) by altering its interactions with specific proteins; 3) by placing the diazirine in a different region of the membrane, crosslinking a different subset of membrane-interacting proteins; and 4) by altering the chemical environment of the diazirine and thus its reactivity.

**Scheme 1.**
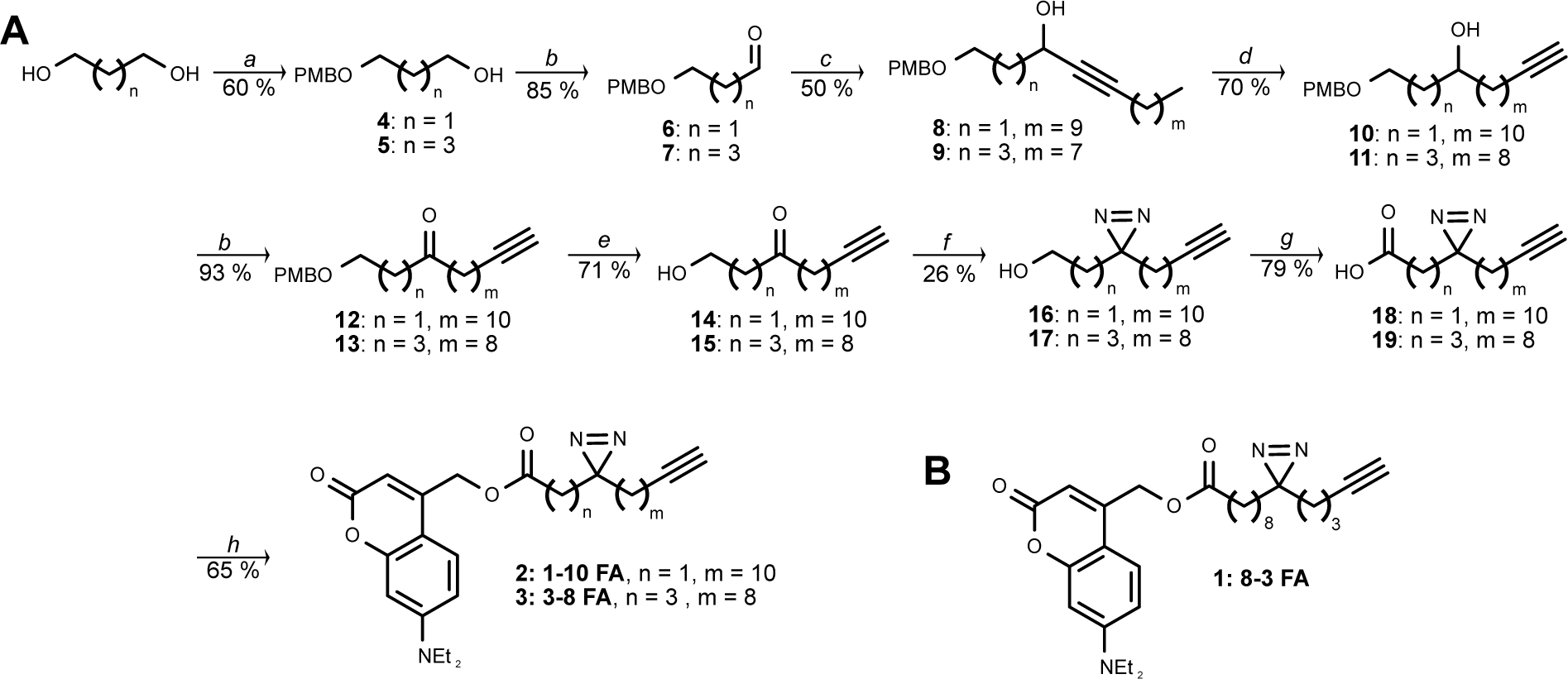
Synthesis of trifunctional fatty acids. (**A**) Synthesis of 1-10 FA and 3-8 FA (*a*) *t*-BuOK, PMBCl, TBAI, THF, 0 °C to rt, with 1,3-propanediol for 1-10 FA and 1,5-pentanediol for 3-8 FA; (*b*) (COCl)_2_, DMSO, then TEA, in DCM, - 85 °C; (*c*) 1-dodecyne (for 1-10 FA) or 1-decyne (for 3-8 FA), *n*-BuLi, THF, -78 °C; (*d*) NaH, diaminopropane, 0 °C; (*e*) DDQ, water/DCM, 0 °C; (*f*) NH_3_, hydroxylaminesulfonic acid, pressure; then I_2_, TEA, methanol, 0°C; (*g*) Jones reagent, acetone, 0 °C; (*h*) 4-methylenehydroxy-7-diethylamino coumarin, DMAP, EDC, DCM, 0 °C to rt (**B**) Structure of 8-3 FA, synthesized as previously described^7^.

In addition to the previously described trifunctional palmitic acid analog with the diazirine at the 10-position, we synthesized two new palmitic acid derivatives with the diazirine at the 3-position and at the 5-position on the acyl chain. We investigated the metabolism and subcellular localization of all probes and showed that the diazirine placement can influence downstream metabolites in different cell lines. We also performed LC-MS/MS-based identification of fatty acid interacting proteins with both distinct and overlapping interactions for each fatty acid derivative. These results provide important considerations for the design of photo-crosslinkable lipid derivatives bearing fatty acids that tolerate the incorporation of a diazirine with minimal effect on biological function.

To synthesize the acyl chains with different diazirine locations, we sought a modular strategy where precursors with any number of carbons could be brought together to produce a diazirine at any desired position on the acyl chain (Scheme 1A). The backbone of the previously synthesized fatty acid (8-3 FA, Scheme 1B, **1**) was prepared based on a Grignard reaction with commercially available TMS-protected chloropentyne^12^. Inserting the diazirine at a different location required a new synthetic strategy. For the 1-10 fatty acid (1-10 FA, **2**) and 3-8 fatty acid (3-8 FA, **3**), one portion of the acyl chain originated from a symmetric diol: symmetrical 1,3-propanediol in the case of 1-10 FA, and 1,5-propanediol in the case of 3-8 FA. The diol was asymmetrically protected with a PMB group, yielding alcohols **4** and **5**, while the other alcohol group was oxidized to aldehydes **6** and **7** by Swern oxidation. The second portion of the final acyl chain was derived from a long-chain terminal alkyne, 1-dodecyne in the case of 1-10 FA and 1-decyne in the case of 3-8 FA. This alkyne was deprotonated by *n*-BuLi to allow it to attack the aldehyde electrophile in the key carbon-carbon bond-forming step to give alcohols **8** and **9**. The well-described “zipper” reaction^13-14^ was employed to isomerize the resulting internal alkyne to the end of the acyl chain, yielding **10** and **11**. After another Swern oxidation, ketones **12** and **13** were deprotected by addition of DDQ, producing **14** and **15**. Next, we performed the ammonia-based and notoriously low-yielding diazirine formation, giving **16** and **17**. After Jones oxidation to bifunctional fatty acids **18** and **19**, we introduced the coumarin cage by a standard esterification reaction to give the caged 1-10 FA derivative **2** and the caged 3-8 FA derivative **3** (Scheme 1A). 8-3 FA (**1**) was synthesized as previously described^7^ and used as the basis of comparison for the novel multifunctional lipid derivatives.

To compare how the fatty acid probes are recognized by their endogenous metabolic enzymes, we performed metabolic labeling with each of the fatty acids to assess which types of lipids they are incorporated into. Previous studies with alkyne-derivatized palmitic acid, as well as radiolabeled palmitic acid^2, 15^, have indicated that the major metabolites of free palmitic acid are phosphatidylcholine (PC), triacylglycerol (TAG), and, to a lesser extent, phosphatidylethanolamine (PE). We treated Huh7 cells with 8-3 FA, 3-8 FA, and 1-10 FA, and extracted the lipids 1 or 24 h after uncaging. Lipid extracts were subjected to click reactions with 3-azido-7-diethylamino-coumarin and separated by TLC (Fig 1: 24 h after uncaging; and Supplementary Figure 1: 1 h after uncaging). After both time points, all three fatty acids were incorporated into PC and TAG, albeit with different distributions. After 24 h of metabolism, we observed some PE accumulation for 8-3 FA and 3-8 FA, but little to none for 1-10 FA. In addition, 1-10 FA showed up much less prominently in TAG after 24 h, although 1-10 FA initially incorporated well into TAG. These results indicate that all three fatty acid backbones are incorporated into the same major subsets of glycerolipids. However, 1-10 FA produced a different metabolic spectrum probably due to subtle differences in the preference of metabolic enzymes for certain diazirine positions.

**Figure 1.**
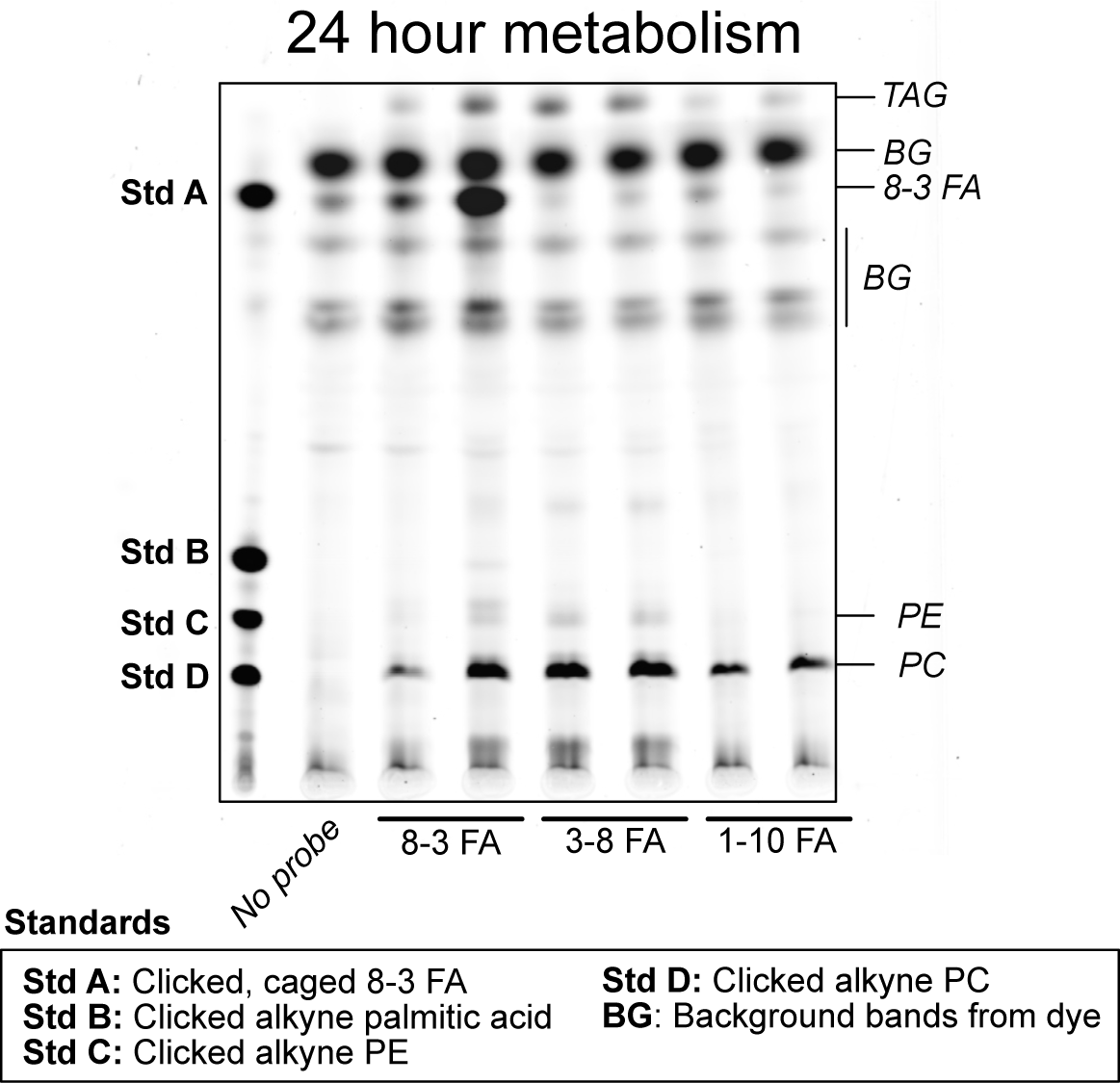
TLC of the three fatty acid probes after 24 h of metabolism in Huh7 cells, 65:25:4 chloroform : methanol : saturated ammonium hydroxide, for 6cm, followed by 1:1 hexanes : ethyl acetate for 9cm.

Next, we wanted to examine the subcellular localization of each lipid probe over time. Bifunctional (uncaged) 8-3 FA was previously described as broadly distributed in membrane-bound organelles^12^. We performed a short time course in the liver-derived Huh7 cells, crosslinking each probe for either 5, 30, or 60 min after uncaging, and co-staining with antibodies against protein disulfide isomerase (PDI) to mark the ER and giantin to mark the Golgi, respectively (Fig 2 and Supplementary Figure 2A). We calculated Pearson’s correlation coefficients (PCC) as a measure of co-localization between the organelle marker and the lipid signal for each cell, using a Cellprofiler pipeline^16-17^. We observed that all three fatty acid probes had relatively even distribution between the ER and Golgi, with PCC between 0.25 and 0.5 for both organelles for all three fatty acids. We did not observe a significant change in probe localization over time for any fatty acid, although 3-8 seems to have preferred an ER location after 60 min (Supplemental Figure 2B). This is in line with the metabolic study described above, as the three fatty acids are incorporated into the same lipid classes, which would be expected to be distributed similarly throughout the cell. This furthermore indicates that the diazirine does not substantially alter the interactions with lipid transport proteins that might transport the fatty acids independent of metabolism.

**Figure 2.**
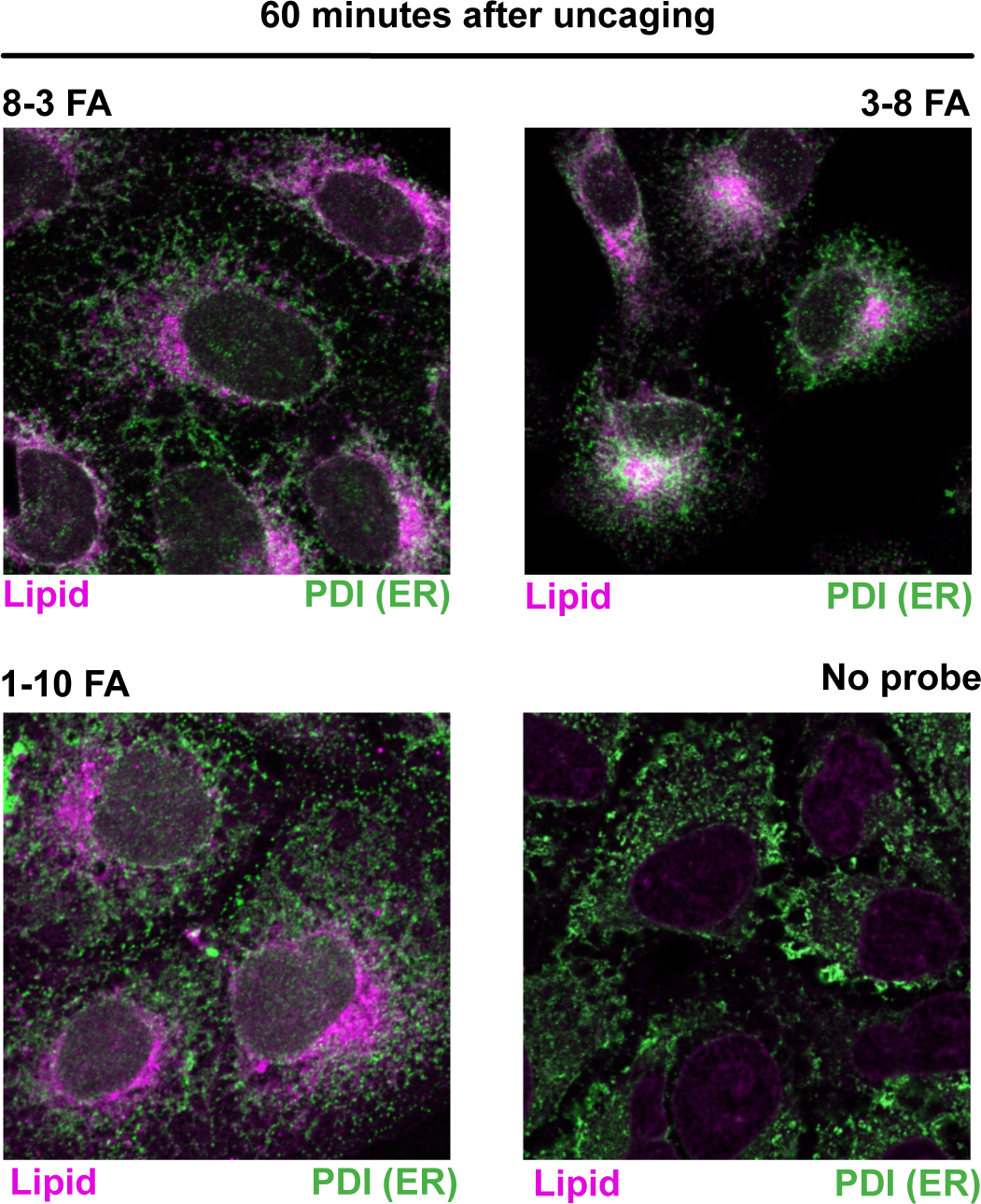
Confocal microscopy of each lipid probe 60 min after metabolism onset in Huh7 cells, after fixation and click chemistry, relative to ER staining. Zeiss LSM 980 Laser-Scanning 4-channel confocal microscope with Airyscan.2.

With the metabolism and localization of each probe established, we next sought to use the photo-crosslinking function of the trifunctional lipid derivatives to compare the protein interactome of each lipid backbone. In-gel fluorescence (Supplementary Figure 5A an 5B) of all three probes shows many similar bands showing up among all three fatty acid probes, although some bands appear exclusively in one probe, and others appear with different intensities among the three. To characterize the interactomes of the fatty acids, we treated Huh7 cells with 8-3 FA and 1-10 FA, representing the two extremes of diazirine position, in two biological replicates, and also included two biological replicates for each probe that had not been exposed to 350 nm light for crosslinking as a negative control. We photo-crosslinked the fatty acid probes 1 h after uncaging and used azide-agarose beads to enrich proteins that had been crosslinked to the alkyne-bearing lipids. We digested off the isolated proteins from the beads with trypsin, as previously described^10^. Peptides were isotopically labeled using the TMT-16 platform for multiplexing and analyzed by LC-MS/MS. Raw signal intensities for each channel were normalized based on variance stabilization^18^, and 1129 proteins were identified. To assess whether there was appreciable enrichment of proteins in the +UV condition over the -UV condition, the ratio of the intensity of each protein in the +UV condition versus the -UV condition was calculated; the log_2_ of this ratio is displayed in Supplementary Figure 3A, highlighting the enrichment of distinct subsets of proteins in each probe condition, and good overall enrichment of proteins for the irradiated conditions.

To identify the most robust interacting proteins for each probe, the normalized signal intensities were subjected to Limma analysis to calculate fold changes and p-values of the intensity in the +UV over the -UV samples (Volcano plots of this analysis are shown in Supplementary Figures 3B and 3C). Hits and candidates were selected based on fold change and false discovery rate criteria: “hit” proteins are those with a fold change greater than 2 and a false discovery rate less than 0.05; “candidate” proteins are those with a fold change greater than 1.5 and a false discovery rate less than 0.2. Three hits and three candidates were identified for 8-3 FA; 22 hits and 10 candidates were identified for 1-10 FA (normalized intensities for all hit and candidate proteins are shown in Supplementary Figures 3D and 3E). A range of proteins were identified for each probe, including many known lipid binding proteins. Several of the hits for 8-3 FA were identified in the previous screen of bifunctional fatty acid, including the fatty acid catabolic enzymes ECH1 and ECHS1 and the phospholipid transport protein PITPNB^12^. 1-10 FA had nearly five times as many hits and candidates as 8-3 FA. While some functionally similar proteins were observed (such as fatty acid catabolic enzymes ACAD9 and ACAA2), there were many more ER resident proteins involved in protein quality control (such as CALML5, UBB, and PDIA6). Only the highly expressing VDAC2 was identified as a hit for both 8-3 FA and 1-10 FA (Figure 3). We selected one candidate protein, PITPNB, to validate by pulldown and western blot (see Supplementary Methods, and Supplementary Figures 5C and 5D). The pulldown recapitulated our findings by proteomics, where 8-3 FA is able to pull down PITPNB much more strongly than 1-10 FA, and further indicated that 3-8 FA had a phenotype more similar to 8-3 FA than 1-10 FA. Overall, these results indicate an overlapping but nonidentical set of protein interactors for the fatty acids with diazirines at either extreme of the acyl chain, despite their structural and metabolic similarities, and their similar subcellular localizations, indicating that the position of the diazirine group will affect the spectrum of targetable proteins.

**Figure 3.**
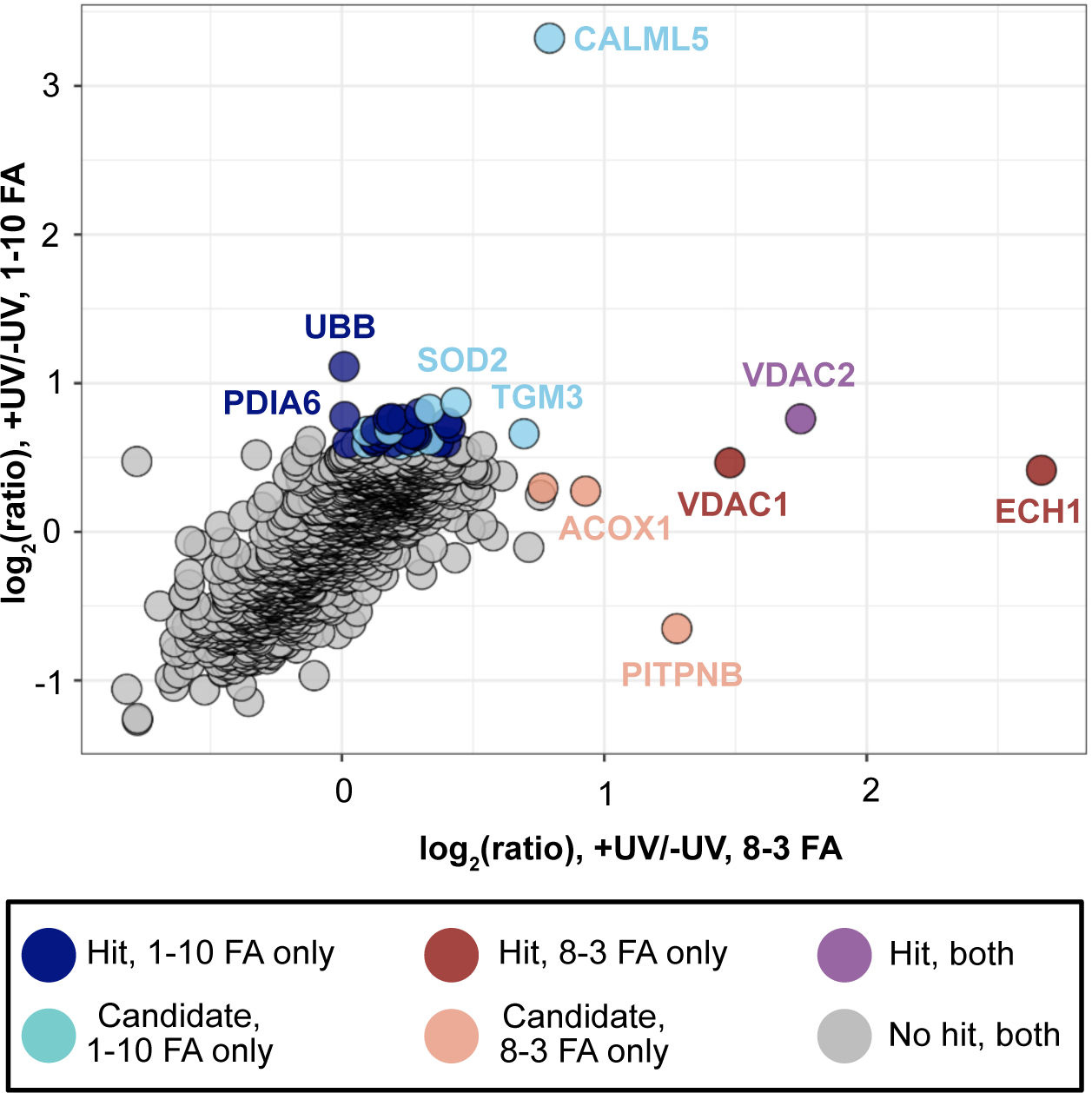
Proteins pulled down by 8-3 FA and 1-10 FA, colored by the degree of enrichment in the (+) UV condition over the (-) UV condition. See text for details.

## CONCLUSIONS

In this work, we present the synthesis of two novel multifunctional lipid probes to profile the interactions of the metabolites of glycerolipids to systematically interrogate how the positioning of the diazirine affects the metabolism, localization, and protein interactions of fatty acid probes. We found that all three fatty acids are converted into a similar pool of glycerolipids, predominantly phosphatidylcholine, with lesser amounts of triacylglycerol, and, after 24 h, phosphatidylethanolamine. We showed that the fatty acids were associated with major membrane-bound organelles, the ER and the Golgi, to similar degrees and in a relatively static fashion over time. Despite their similarity in metabolism and subcellular localization, we observed striking differences between the interacting partners of the two fatty acid probes 1-10 FA and 8-3 FA. Interestingly, only one protein directly overlapped between the hits from both fatty acids (the mitochondrial scramblase VDAC2). There was some functional overlap between proteins of fatty acid metabolism (especially in the beta oxidation pathway; ECH1 and ECHS1 for 8-3 FA versus ACAA2 and ACAD9 for 1-10 FA). 1-10 FA in general had many more interacting partners, either due to its position making it more accessible to peripheral membrane proteins, or because the different chemical environment affected the inherent reactivity of the respective diazirines, as probe structure is known to have a profound effect on the photo-crosslinking efficiency of a diazirine^19^. This echoes experiments with bifunnctional myristic acid derivatives, which also found more protein interactors for probes with the diazirine closer to the carboxylic acid ^20^. These findings underscore the importance of diazirine placement in the design of lipid photoaffinity probes, as the same base molecule can yield dramatically different labeling profiles based solely on the location of the diazirine.

## ACKNOWLEDGMENTS

This work was supported by funding from the National Institutes of Health (NIAID R01 AI141549-02, to FGT and CS and R01 GM127631 to CS) and an endowment to CS, donated by Helen Jo and Bill Whitsell.

## METHODS

### Synthesis

All chemicals were purchased from commercial suppliers and were used without further purification. Solvents were of ACS chemical grade (Fisher Scientific) and were used without further purification. Analytical thin-layer chromatography was performed on silica gel 60 F254 aluminum-backed plates (Millipore Sigma) and spots were visualized either by UV light (254 nm for PMB-containing compounds; 365 nm for coumarin-containing compounsd) or potassium permanganate staining (1.5 g KMnO_4_, 10g K_2_CO_3_, and 1.25 mL 10% NaOH in 200 mL of water). Flash column chromatography was performed with manually packed columns using Thermo Scientific Chemical Silica gel (0.035-0.070mm, 60 Å). High pressure liquid chromatography (HPLC) was performed on a Varian Prostar 210 (Agilent) using Polars 5 C18-A columns (Analytical: 150 x 4.6 mm, 3 µm; Preparative: 150 x 21.2 mm, 5 µm). Mass spectra were obtained on an Advion Expressions CMS mass spectrometer. ^1^H NMR spectra were recorded on a Bruker DPX spectrometer at 400 MHz. Chemical shifts are reported as parts per million (ppm) downfield from solvent references. Spectral characterization can be found in the Supporting Information.

#### Synthesis of 1-10 FA (2)

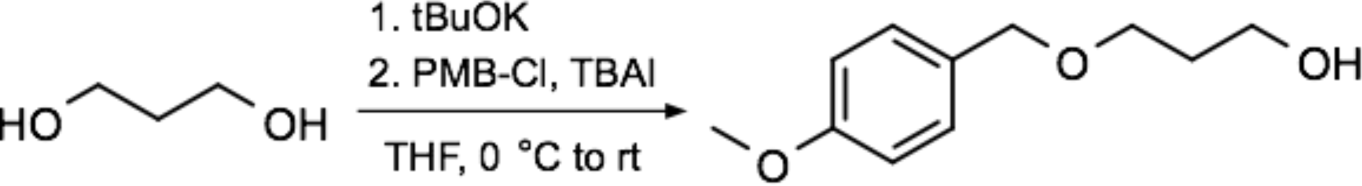

**3-[4-Methoxyphenyl)methoxy]propan-1-ol (4).** To a 0°C solution of 1,3-propanediol (18.9 mL, 262.8 mmol, 1 equivalent) in anhydrous THF (150 mL) was added a suspension of potassium *tert*-butoxide (14.75 g, 131 mmol, 1 equivalent) in anhydrous THF (150 mL), dropwise. To the resulting viscous suspension of alkoxide was added tetrabutylammonium iodide (9.7083 g, 18.95 mmol, 0.075 equivalents) and para-methoxylbenzyl chloride (14.4 mL, 0.5 equivalents). Once the addition was complete, the reaction was allowed to warm to room temperature and stirred overnight. TLC (2:1 hexanes : ethyl acetate) showed the formation of the desired product (Rf = 0.2) as well as the bis-protected byproduct (Rf = 0.75). Water (100 mL) was added, and the aqueous layer was extracted 3 x 100 mL with ethyl acetate and the combined organic layers were washed with brine (200 mL), dried over MgSO_4_, and concentrated under reduced pressure. The orange oil was purified by flash chromatography, with a gradient of 2 : 1 hexanes : ethyl acetate to pure ethyl acetate, yielding **4**, 15.3 g, in 60% yield. ^1^H NMR (400 MHz, CDCl_3_) δ = 7.255 (dt, J = 8.8, J = 2.8, 2H), 6.881(dt, J = 8.8, J = 2.8, 2H), 4.455(s, 2H), 3.806(s, 3H), 3.776(t, J = 5.6, 2H), 3.641(t, J = 5.6, 2H), 1.855(quint, J = 5.6, 2H); ^13^C NMR (400 MHz, CDCl_3_) δ = 159.2, 130.2, 129.3, 113.8, 68.9, 61.7, 55.3, 32.1; MS (*m/z*): [M - H_2_O + H]^+^ calcd. for C_11_H_16_O_3_, 179.107; found 179.2

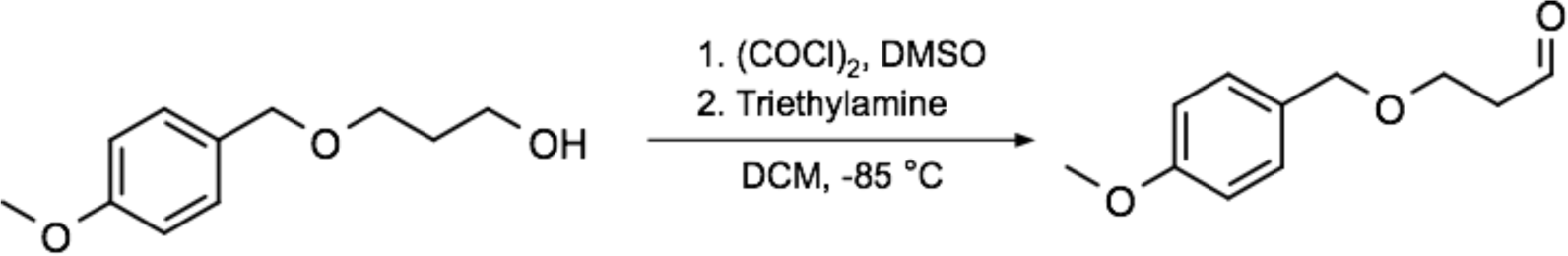

**3-[4-Methoxyphenyl)methoxy]propanal (6).** Dimethyl sulfoxide (10.8 mL, 152.6 mmol, 2.4 equivalents) was diluted in dichloromethane (10 mL) and added, dropwise, to a stirring solution of oxalyl chloride (6 mL, 69.95 mmol,, 1.1 equivalents) in dichloromethane (200 mL) at -85°C. After the evolution of gas had ceased, **4** (12.48 g, 63.59 mmol, 1 equivalent), diluted in 10 mL dichloromethane, was added dropwise. This mixture was stirred at -85°C for thirty min, when the appearance of the sulfonium intermediate by TLC (3:1 hexanes : ethyl acetate, Rf = 0), and then triethylamine (40 mL, 292.5 mmol, 4.6 equivalents) was added, dropwise, and product was observed by TLC (3:1 hexanes : ethyl acetate, Rf = 0.6). The reaction was allowed to warm to room temperature, and then 200 mL of water was added. The aqueous layer was extracted 2x100mL with dichloromethane. The combined organic layers were washed 2 x 200 mL with 10% citric acid, 1 x 100 mL brine, then dried over MgSO_4_ and concentrated under reduced pressure, yielding 11.56 g of **6** (85 % yield). ^1^H NMR (400 MHz, CDCl_3_) δ = 9.786(t, J = 1.8, 1H), 7.26(dt, J = 8.8, J = 2.8, 2H), 6.891(dt, J = 8.8, J = 8.8, 2H), 4.463(s, 2H), 3.805(s, 3H), 3.785(t, J = 6, 2H), 2.684(td, J =, J = 2, 2H); ^13^C NMR (400 MHz, CDCl_3_) δ = 201.3, 159.4, 130.0, 129.5, 113.9, 73.0, 63.6, 55.4, 44.0

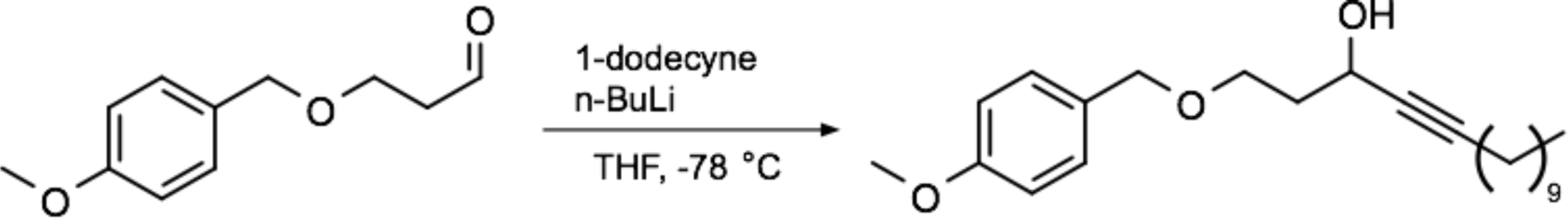

**1-[(4-Methoxyphenyl)methoxy]pentadec-4-yn-3-ol (8).** To a -78 °C solution of 1-dodecyne (11.5 mL, 54.01 mmol, 1 equivalent) in anhydrous tetrahydrofuran (200 mL) was added 2.5 M *n*- butyllithium (32.4 mL, 81.01 mmol, 1.5 equivalents), dropwise. This was stirred for thirty min, producing an off-white slurry, following which a 0 °C solution of **6** (10.49 g, 54.01 mmol, 1 equivalent) in anhydrous tetrahydrofuran (100 mL) was added, dropwise. The reaction was stirred for 1 h at -23 °C, then overnight at room temperature. Following the appearance of product by TLC (4:1 hexanes : ethyl acetate, Rf = 0.5), the reaction was quenched with 10 % citric acid and then acidified to pH 4 with the same. A further 200 mL water was added, and then the aqueous layer was extracted 2 x 150 mL with ethyl acetate, washed 1 x 200 mL with brine, dried over magnesium sulfate and concentrated under reduced pressure. The crude reside was purified by flash chromatography (4:1 hexanes : ethyl acetate), yielding 19.67 g of **8** as a yellow oil (50.6 % yield). ^1^H NMR (400 MHz, CDCl_3_) δ = 7.267(dt, J = 8.8, J = 2.8, 2H), 6.908(dt, J = 8.8, J = 2.8, 2H), 4.596(m, 1H), 4.482(s, 2H), 3.823(m, 1H), 3.803(s, 3H), 3.662(m, 1H), 2.210(td, J = 7, J = 2, 2H), 2.066(m, 1H), 1.943(m, 1H), 1.577(br, 1H), 1.509(quint, J = 7, 4H), 1.380(m, 2H), 1.282(s, 12H), 0.916(t, J = 6.8, 2H); MS (*m/z*): [M - H_2_O + H]^+^ calcd. for C_23_H_36_O_3_, 343.263; found 343.5

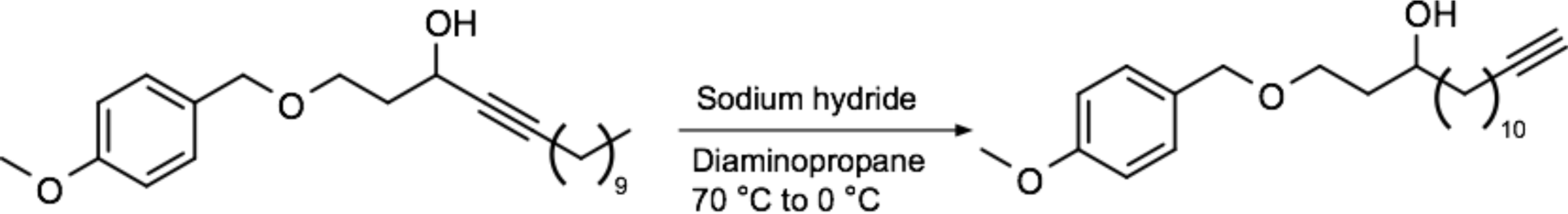

**1-[4-Methoxyphenyl)methoxy]pentadec-14-yn-3-ol (10).** 60% sodium hydride in mineral oil (3.003 g, 75.02 mmol, 5.5 equivalents) was suspended in diaminopropane (50 mL) under argon atmosphere. The suspension was heated to 70 °C and stirred for 1 h, resulting in a clear brown solution. This solution was cooled to 0 °C and **8** (4.917 g, 13.64 mmol, 1 equivalent) was added, dropwise, resulting in a dark, red-black solution. Once TLC showed consumption of starting material, after about 15 min (4:1 hexanes : ethyl acetate, product Rf = 0.45), the reaction was quenched by pouring slowly over a large excess of ice, extracted 3 x 50 mL with dichloromethane, and the combined organic layers were washed 2 x 50 mL with 10% ice-cold citric acid and 1 x 50 mL brine, dried over magnesium sulfate, and concentrated. The crude orange residue was purified by flash chromatography (4:1 hexanes : ethyl acetate), yielding an orange oil, **10**, 3.48 g (70.8 %). ^1^H NMR (400 MHz, CDCl_3_) δ = 7.26(dt, J = 8.8, J = 2.8, 2H), 6.884(dt, J = 8.8, J = 2.8, 2H), 4.452(s, 2H), 3.803(s, 3H), 3.684(m, 1H), 3.626(m, 1H), 2.173(td, J = 7, J = 2.4, 2H), 1.835(t, J = 2.4, 1H), 1.714(m, 2H), 1.538(m, 6H), 1.397(m, 5H), 1.271(s, 12H); ^13^C NMR (400 MHz, CDCl_3_) δ = 159.4, 130.1, 129.4, 113.9, 73.1, 71.7, 69.2, 68.1, 55.4, 37.6, 36.5, 29.8, 29.7, 29.6, 29.6, 29.2, 28.9, 28.6, 27.9, 25.7, 18.5; MS (*m/z*): [M - H_2_O + H]^+^ calcd. for C_23_H_36_O_3_, 343.263; found 343.5

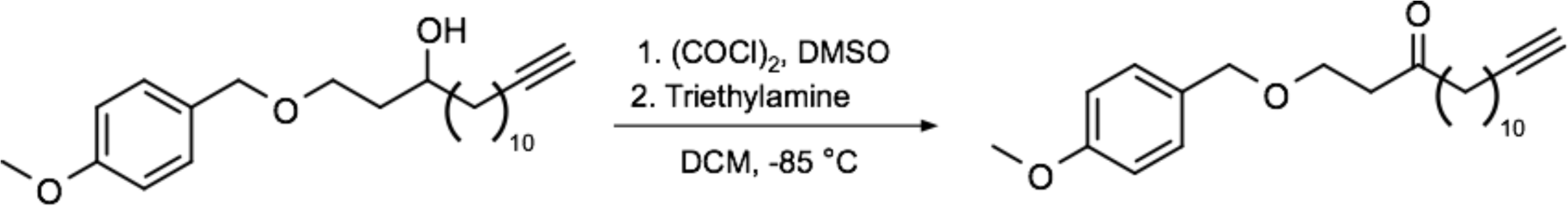

**1-[4-Methoxyphenyl)methoxy]pentadec-14-yn-3-one (12).** Dimethyl sulfoxide (4.4 mL, 62.34 mmol, 2.4 equivalents) was diluted in dichloromethane (5 mL) and added, dropwise, to a stirring solution of oxalyl chloride (2.5 mL, 28.573 mmol, 1.1 equivalents) in dichloromethane (200 mL) at -85°C. After the evolution of gas had ceased, **10** (9.365 g, 25.975 mmol, 1 equivalent) was added dropwise. This mixture was stirred at -85°C for thirty min, and then triethylamine (16.7 mL, 119.49 mmol, 4.6 equivalents) was added, dropwise. The reaction was allowed to warm to room temperature over 1 h, and after the appearance of product by TLC (4:1 hexanes : ethyl acetate, Rf = 0.6), 200 mL of water were added. The aqueous layer was extracted 2x100mL with dichloromethane. The combined organic layers were washed 2 x 200 mL with 10% citric acid, 1 x 100 mL brine, then dried over MgSO_4_ and concentrated under reduced pressure, yielding 8.635 g of **12** (93% yield) as a yellow solid. ^1^H NMR (400 MHz, CDCl_3_) δ = 7.223(dt, J = 8.8, J = 2.8, 2H), 6.877(dt, J = 8.8, J = 2.8, 2H), 4.430(s, 2H), 3.797(s, 3H), 3.702(t, J = 6.4, 2H), 2.668(t, J = 6.4, 2H), 2.418(t, J = 7.6, 2H), 2.177(td, J = 7, J = 2.4, 2H), 1.933(t, J = 2.4, 1H), 1.496(m, 4H), 1.375(m, 2H), 1.258(s, 12H); ^13^C NMR (400 MHz, CDCl_3_) δ = 209.7, 159.3, 130.3, 129.4, 113.9, 84.9, 73.0, 65.5, 60.5, 55.4, 43.5, 43.2, 42.9, 29.5, 29.5, 29.3, 29.2, 28.8, 28.6, 25.8, 23.7, 2.4, 21.1, 18.5, 14.3, 13.9; MS (*m/z*): [M + H]^+^ calcd. for C_23_H_34_O_3_, 359.258; found 359.3

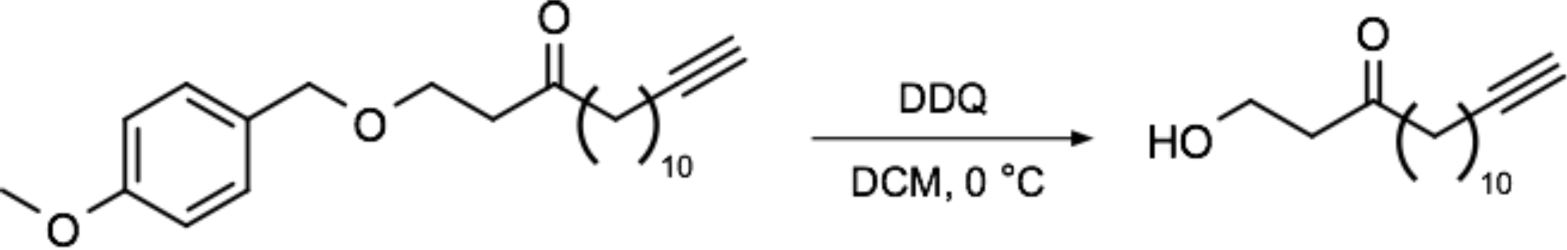

**1-Hydroxypentadec-14-yn-3-one (14).** To a 0 °C solution of **12** (8.635 g, 24.08 mmol, 1 equivalent) in DCM (500 mL), 2,3-dichloro-5,6-dicyano-1,4-benzoquinone was added as a solid, followed by water (50 mL). The reaction was stirred for 30 min at 0 °C and then for 1 h at room temperature. Once the reaction had gone to completion, by TLC (4:1 hexanes : ethyl acetate, Rf = 0.1, KMnO_4_ staining), the reaction was poured over 500 mL saturated sodium bicarbonate, extracted 2 x 200 mL DCM, then washed 2 x 300 mL with 10% sodium bicarbonate and 2 x 300 mL brine. The crude orange residue was purified by flash chromatography, 3:1 hexanes:ethyl acetate, yielding **14** (4.3 g, 17.17 mmol) as a pale orange semi-crystalline solid in 70.8 % yield. ^1^H NMR (400 MHz, CDCl_3_) δ = 3.837(m, 2H), 2.666(t, J = 5.4, 2H), 2.431(t, J = 7.4, 2H), 2.175(td, J = 7, J = 2.4, 2H), 1.937(t, J = 2.4, 1H), 1.499(m, 4H), 1.378(m, 2H), 1.275(s, 10H); ^13^C NMR (400 MHz, CDCl_3_) δ = 68.1, 57.7, 35.7, 33.3, 29.5, 29.4, 29.2, 29.1, 29.0, 28.8, 28.5, 23.9, 18.5

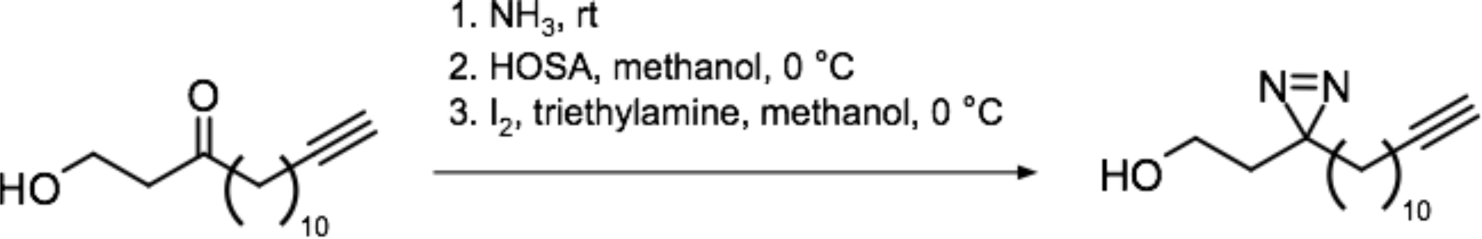

**2-[3-(Dodec-11-yn-1-yl)-3H-diazirin-3-yl]ethan-1-ol (16). 14** (3.5 g, 1 equivalent) was placed in an oven-dried bomb flask and cooled to -85 °C. Ammonia (25 mL) was condensed on top of the compound and the bomb flask was capped. The mixture was allowed to warm to room temperature, stirring, whereupon **14** dissolved in the ammonia. The reaction stirred for 24hr at room temperature, and then the ammonia was evaporated overnight, into an aqueous 10% citric acid bath. Hydroxylaminesulfonic acid (3.65 g, 2.2 equivalents) was dissolved in 15 mL anhydrous methanol and added to the residue in the bomb flask at 0 °C. This was stirred for 1 h at rt, and then the reaction mixture was filtered to remove white solids and the methanol was evaporated. The residue was dissolved in 50 mL ethyl acetate and washed 1x 25 mL 10% citric acid and 1x25 mL brine, dried over MgSO4, and concentrated. The crude residue was oxidized without further purification, by dissolving in 50 mL methanol, cooling to 0 °C, adding TEA (4.1 mL, 2 equivalents), and I_2_ (3.73 g, 1 equivalent) portion wise as a solid until the brown iodine color persisted. Methanol was once again evaporated, the residue was dissolved in 50 mL ethyl acetate and washed 4x 50mL with sodium thiosulfate and 1x 50mL brine. TLC (4:1 hexanes : ethyl acetate) indicated consumption of starting material and a complex mixture of products; isolation of the spot with Rf = 0.65 resulted in the correct product. The crude reside was purified by flash chromatography (4:1 hexanes : ethyl acetate) yielding **16** (0.939 g) in 25.6 % yield as a pale yellow solid. ^1^H NMR (400 MHz, CDCl_3_) δ = 3.454(m, 2H), 2.171(td, J = 7, J = 2.4, 2H), 1.937(t, J = 2.4, 1H), 1.679(t, J = 6.4, 2H), 1.517(m, 2H), 1.413(m, 4H), 1.253(m, 15H), 1.090(m, 3H); MS (*m/z*): [M + NH4]^+^ calcd. for C_15_H_26_N_2_O, 268.238; found 268.1

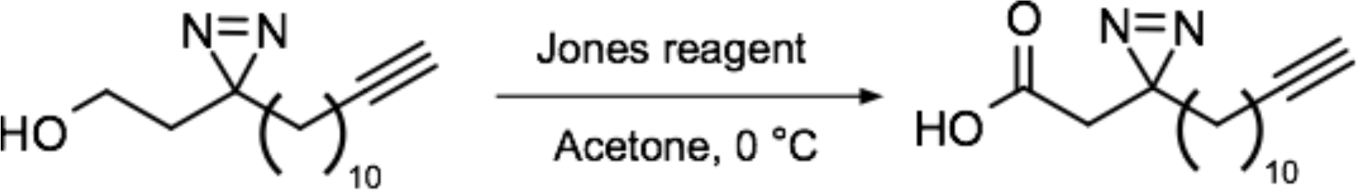

**2**-**[3**-**(Dodec**-**11**-**yn**-**1**-**yl)**-**3H**-**diazirin**-**3**-**yl]acetic acid (18)**. **16** (50 mg, 1 equivalent) was dissolved in acetone (20 mL) and cooled to 0 °C. Jones reagent (2 M, 100µL, 1 equivalent) was added, dropwise, until the pink color of the reagent persisted. The reaction was stirred for 15 min at 0°C, and after the appearance of product by TLC (8:1 hexanes : ethyl acetate, Rf = 0.05, KMnO_4_ staining) was quenched with isopropanol, producing a bright, blue-green precipitate. The precipitate was filtered and then the acetone was removed under reduced pressure. The residue was dissolved in ethyl acetate (20 mL) and washed twice with saturated sodium bicarbonate and once with brine, dried over magnesium sulfate, and concentrated, yielding pure **18** (41.9 mg, 79.3 %) as a white solid. ^1^H NMR (400 MHz, CDCl_3_) δ = 2.301(s, 2H), 2.175(td, J = 7, J = 2.4, 2H), 1.936(t, J = 2.4, 1H), 1.536(m, 5H), 1.379(m, 3H), 1.259(m, 12H), 1.086(m, 3H); MS (*m/z*): [M + 2H]^+^, calcd. for 2(C_15_H_24_N_2_O_2_), 530.382, found 530.3.

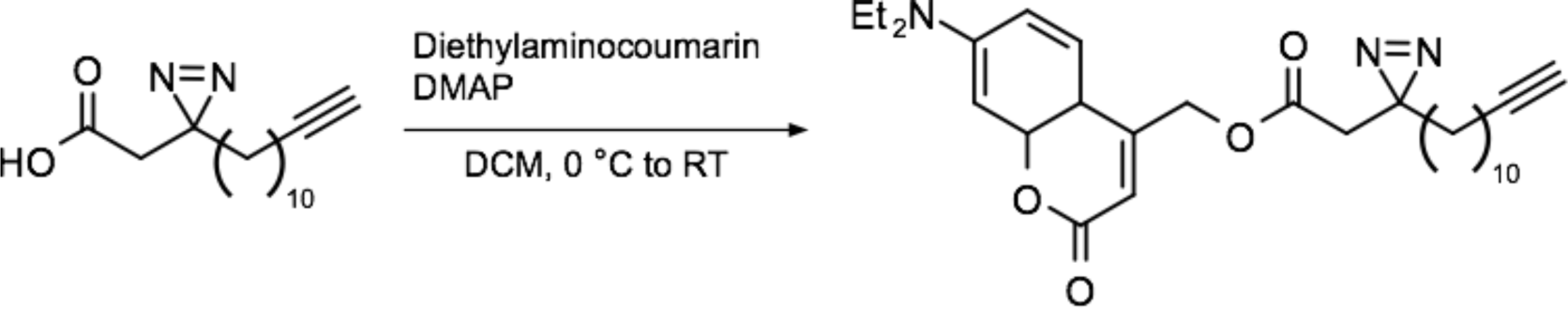

**[7-(Diethlamino)-2-oxo-2H-chromen-4-yl]methyl 2-[3-(prop-2-yn-1-yl)-3H-diazirin-3-yl] acetate (1-10 FA). 8** (100 mg, 1 equivalent) was dissolved in 10 mL DCM. EDC (163 mg, 2.1 equivalents) and DMAP (10 mg, 0.2 equivalents) were added and stirred together for 15 min, as the mixture gradually turned orange. 7-(diethylamino) 4-hydroxymethylcoumarin was dissolved in 5 mL DCM. The coumarin and the activated compound **8** were both cooled to 0 °C and the coumarin added was added to compound **8**. The reaction was allowed to warm to room temperature overnight. Appearance of product was observed by TLC (2:1 hexanes : ethyl acetate, Rf = 0.2), and the reaction was poured over citric acid, extracted 2 x 15 mL with DCM, washed once with brine, and concentrated under reduced pressure. The crude residue was purified initially by flash chromatography (2:1 hexanes : ethyl acetate), and then by preparative LC-MS, with a gradient of 50-100% acetonitrile in water over 35 min; pure product appeared at 18.33 min as a pale yellow solid (130 mg, 64.9 % yield). ^1^H NMR (400 MHz, CDCl_3_) δ = 7.329 (d, J = 8.8, 1H), 6.649 (dd, J = 8.6, J = 1.6, 1H), 6.571 (d, J = 2.8, 1H), 6.192 (t, J = 1.2, 1H), 5.270 (d, J = 1.6, 2H), 3.439 (q, J = 7.2, 4H), 2.400 (s, 2H), 2.191 (td, J = 7.2, J = 2.4, 2H), 1.942 (t, J = 2.4, 2H), 1.536 (m, 6H), 1.382 (m, 2H), 1.243 (m, 22H), 1.083 (m, 2H); ^13^C NMR (400 MHz, CDCl_3_) δ = 168.8, 161.8, 156.4, 150.5, 148.8, 124.6, 109.3, 107.3, 98.5, 84.9, 68.2, 62.1, 45.2, 39.8, 32.7, 29.9, 29.5, 29.5, 29.4, 29.2, 29.2, 28.9, 28.6, 26.2, 23.9, 18.5, 12.5 (m/z): [M + H]+, calcd. For C_29_H_39_N_3_O_4_, 494.2948, found 494.3

#### Synthesis of 3-8 FA (3)

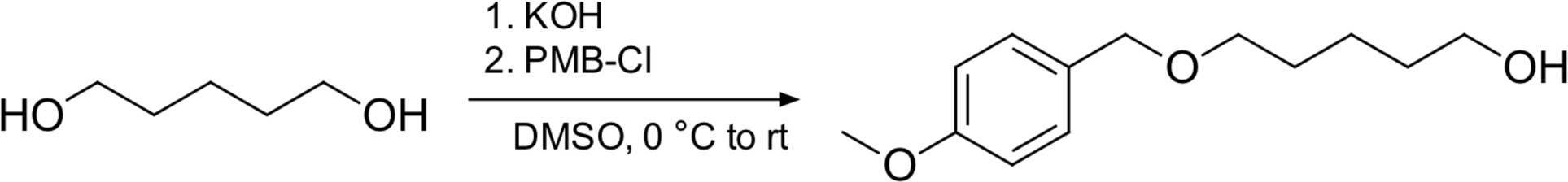

**5-[(4-Methoxyphenyl)methoxy]pentan-1-ol (5)** To a 0 C° solution of 1,5-pentanediol (9.5 g, 91.21 mmol, 1 equivalent) in dimethyl sulfoxide (40 mL) was added potassium hydroxide (8 g, 91.21 mmol, 1 equivalent). The reaction was stirred at room temperature for 30 min, and then cooled to 0 °C for the addition of para-methoxybenzyl chloride (7.142 g, 45.60 mmol, 0.5 equivalents), and then returned to room temperature and stirred for 1.5 h. TLC (2:1 hexanes : ethyl acetate, UV visualization) showed the formation of the desired product (Rf = 0.15) as well as the bis-protected by-product (Rf = 0.6). Water (100 mL) was added, and the aqueous layer was extracted 3 x 100 mL with ether and the combined organic layers were washed with brine (200 mL), dried over MgSO_4_, and concentrated under reduced pressure. The orange oil was purified by flash chromatography, with a gradient of 20% ethyl acetate in hexanes to 50% ethyl acetate in hexanes, yielding **5**, 15.0 g, in 73.3% yield. 1H NMR (400 MHz, CDCl3) δ = 7.25 (d, J = 8.8, 2H), 6.86 (d, J = 8.8, 2H), 4.24 (s, 2H), 3.79 (s, 3H), 3.62 (t, J = 6.4, 2H), 3.44 (t, J = 6.4, 2H), 1.66-1.41 (m, 6H); 13C NMR (400 MHz, CDCl3) δ = 1591.2, 130.7, 129.3, 113.8, 72.6, 70.0, 62.8, 55.3, 32.5, 29.5, 22.5

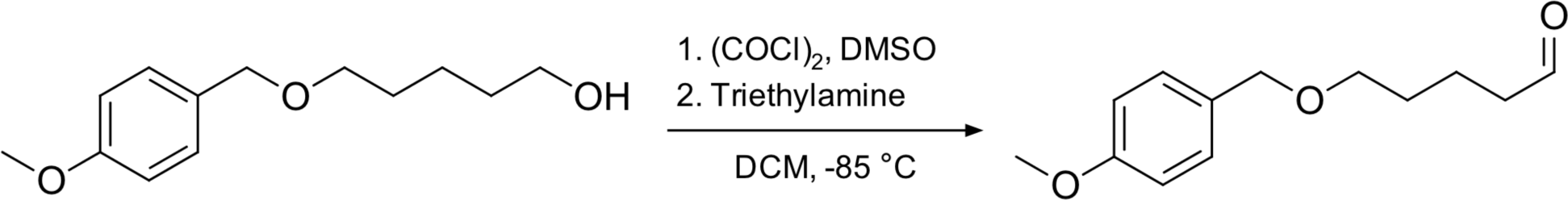

**5-[(4-Methoxyphenyl)methoxy]pentanal (7)** Dimethyl sulfoxide (12.18 mL, 171.32 mmol, 2.4 equivalents) was diluted in dichloromethane (10 mL) and added, dropwise, to a stirring solution of oxalyl chloride (9,97 g, 78.82 mmol,, 1.1 equivalents) in dichloromethane (50 mL) at -85°C. After the evolution of gas had ceased, **5** (16.0 g, 71.38 mmol, 1 equivalent), diluted in 10 mL dichloromethane, was added dropwise. This mixture was stirred at -85°C for thirty min, with the appearance of the sulfonium intermediate by TLC (2:1 hexanes : ethyl acetate, Rf = 0). Triethylamine 45.46 mL, 328.36 mmol, 4.6 equivalents) was then added, dropwise, and product was observed by TLC (2:1 hexanes : ethyl acetate, Rf = 0.4, UV visualization). The reaction was allowed to warm to room temperature, and then 200 mL of water was added. The aqueous layer was extracted 2x100mL with dichloromethane. The combined organic layers were washed 2 x 200 mL with 10% citric acid, 1 x 100 mL brine, then dried over MgSO_4_ and concentrated under reduced pressure, yielding 14.96 g of **7** (67 % yield). ^1^H NMR (400 MHz, CDCl_3_) δ = 9.75 (t, J = 1.6, 1H), 7.25 (d, J = 8.8, 2H), 6.87 (d, J = 8.8, 2H), 4.42 (s, 2H), 3.8-(s, 3H), 3.46 (t, J = 6, 2H), 2.45 (td, J = 7.2, J = 1.6, 2H), 1.77-1.6 (m, 4H); ^13^C NMR (400 MHz, CDCl_3_) δ = 202.6, 159.2, 130.5, 129.3, 113.8, 72.6, 69.5, 55.3, 43.6, 29.1, 18.9

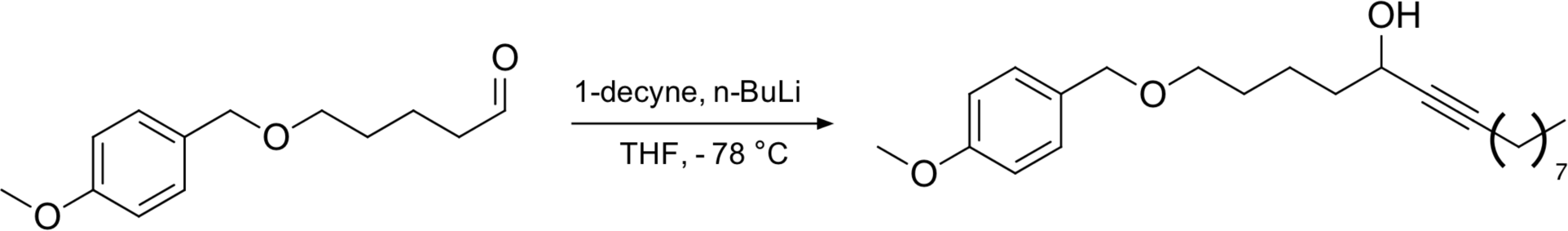

**1-[(4-Methoxyphenyl)methoxy]pentadec-6-yn-5-ol (9)** To a -78 °C solution of 1-decyne (14.04 mL, 94.34 mmol, 1.5 equivalent) in anhydrous tetrahydrofuran (200 mL) was added 2.5 M n-butyllithium (37 mL, 94.54 mmol, 1.5 equivalents), dropwise. This was allowed to warm to 0 °C and stirred for 1h, following which a 0 °C solution of **7** (14.0 g, 63.024 mmol, 1 equivalent) in anhydrous tetrahydrofuran (10 mL) was added, dropwise. The reaction was stirred for 1 h at 0 °C, then overnight at room temperature. Following the appearance of product by TLC (2:1 hexanes : ethyl acetate, Rf = 0.4, UV visualization), the reaction was quenched with 10 % citric acid and then acidified to pH 4 with the same. A further 200 mL water was added, and then the aqueous layer was extracted 2 x 150 mL with ethyl acetate, washed 1 x 200 mL with brine, dried over magnesium sulfate and concentrated under reduced pressure. The crude reside was purified by flash chromatography (gradient of 10 % ethyl acetate in hexanes to 20 % ethyl acetate in hexanes), yielding 17.007 g of **9** as a yellow oil (50.6 % yield). ^1^H NMR (400 MHz, CDCl_3_) δ = 7.26 (d, J = 8.8, 2H), 6.87 (d, J = 8.8, 2H), 4.43 (s, 2H), 4.36 (t, J = 7.2, 1H), 3.8 (s, 3H), 3.44 (t, J = 6, 2H), 2.19 (td, J = 7.2, J = 1.6, 2H), 1.72-1.61 (m, 4H), 1.56-1.46 (m, 4H), 1.27 (m, 10H), 0.88 (t, U = 7.2, 2H); ^13^C NMR (400 MHz, CDCl_3_) δ = 159.1, 130.7, 129.2, 113.8, 85.7, 81.2, 72.6, 70.0, 62.7, 55.3, 38.9, 31.8, 29.4, 29.2, 29.1, 28.9, 28.7, 22.7, 22.0, 18.7, 14.1

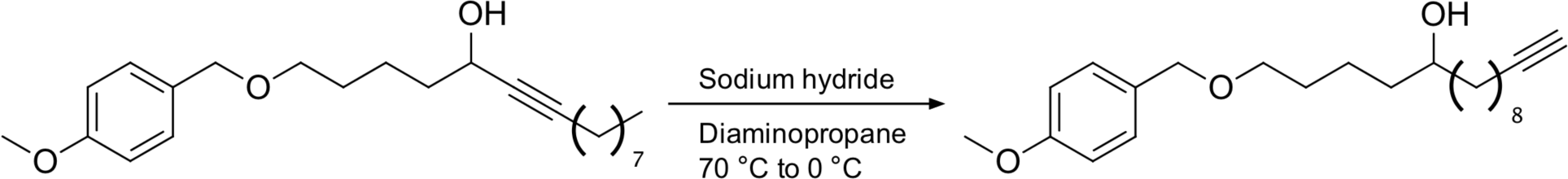

**1-[(4-Methoxyphenyl)methoxy]pentadec-14-yn-5-ol (11)** 60% sodium hydride in mineral oil (1.831 g, 76.33 mmol, 5.5 equivalents) was suspended in diaminopropane (50 mL) under argon atmosphere. The suspension was heated to 70 °C and stirred for 1 h, resulting in a clear brown solution. This solution was cooled to 0 °C and **9** (5 g, 13.82 mmol, 1 equivalent), dissolved in 5 mL diaminopropane was added, dropwise, resulting in a dark, red-black solution. Once TLC showed consumption of starting material, after about 2 h (2:1 hexanes : ethyl acetate, product Rf = 0.3, UV visualization), the reaction was quenched by diluting in 50 mL DCM and slowly adding 100 mL ice-cold water, extracted 3 x 50 mL with dichloromethane, and the combined organic layers were washed 2 x 50 mL with 10% ice-cold citric acid and 1 x 50 mL brine, dried over magnesium sulfate, and concentrated. The crude orange residue was purified by flash chromatography (20 % ethyl acetate in hexanes), yielding an orange oil, **11**, 2.31 g (46.4 %). ^1^H NMR (400 MHz, CDCl_3_) δ = 7.19 (d, J = 8.8, 2H), 6.80 (d, J = 8.8, 2H), 4.36 (s, 2H), 3.74 (s, 3H), 3.51 (m, 1H), 3.38 (t J = 6.4, 2H), 2.11 (td(J = 7.2, J = 2.8, 2H), 1.87 (t, J = 2.8, 1H), 1.53 (m, 4H), 1.47-1.19 (m, 16H); ^13^C NMR (400 MHz, CDCl_3_) δ = 1591., 130.7, 129.3, 113.8, 84.8, 72.6, 71.8, 70.0, 68.1, 55.3, 37.5, 37.2, 29.7, 29.6, 29.5, 29.1, 28.7, 28.5, 25.6, 22.4, 18.4

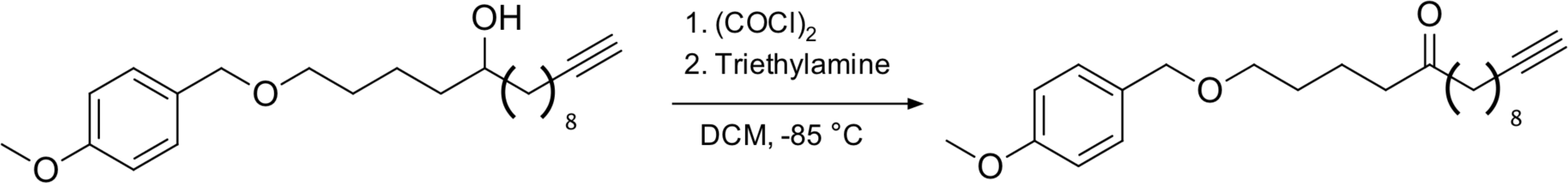

**1-[(4-Methoxyphenyl)methoxy]pentadec-14-yn-5-one (13)** Dimethyl sulfoxide (0.85 mL, 4.99 mmol, 2.4 equivalents) was diluted in dichloromethane (5 mL) and added, dropwise, to a stirring solution of oxalyl chloride (0.47 mL, 5.49 mmol, 1.1 equivalents) in dichloromethane (50 mL) at -85°C. After the evolution of gas had ceased, **11** (1.8 g, 4.99 mmol, 1 equivalent) was added dropwise. This mixture was stirred at -85°C for thirty min, and then triethylamine (3.2 mL, 22.98 mmol, 4.6 equivalents) was added, dropwise. The reaction was allowed to warm to room temperature over 1 h, and 200 mL of water were added. The aqueous layer was extracted 2x100 mL with dichloromethane. The combined organic layers were washed 2 x 200 mL with 10% citric acid, 1 x 100 mL brine, then dried over MgSO_4_ and concentrated under reduced pressure, yielding 1.7 g of **13** (93 % yield) as a yellow solid. ^1^H NMR (400 MHz, CDCl_3_) δ = 7.12 (d, J = 8.8, 2H), 6.81 (d, J = 8.8, 2H), 4.35 (s, 2H), 3.74 (s, 3H), 3.37 (t = J = 6.4, 2H), 2.35 - 2.28 (m, 4H), 2.11 (td, J = 7.2, J = 2.8, 2H), 1.87 (t, J = 2.8, 1H), 1.58-1.41 (m, 8H), 1.32-1.29 (m, 2H), 1.47-1.19 (m, 6H); ^13^C NMR (400 MHz, CDCl_3_) δ = 159.1, 130.7, 129.3, 113.8, 83.4, 76.0, 72.6, 70.1, 70.0, 55.3, 36.0, 31.8, 29.7, 29.0, 28.9, 28.8, 27.8, 22.6, 22.4, 18.8, 14.1

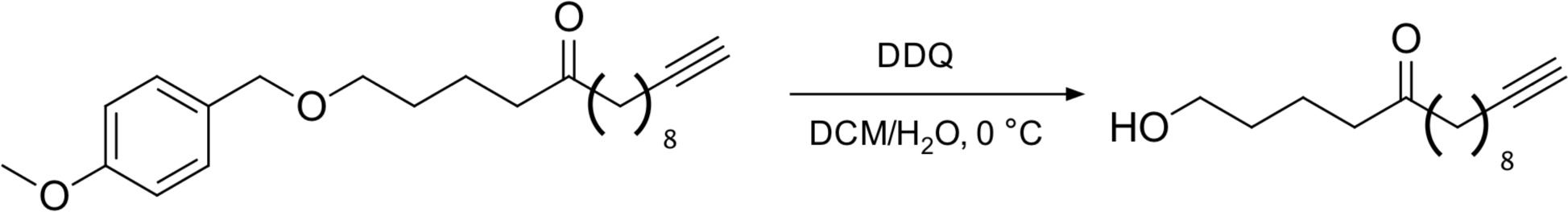

**1-Hydroxypentadec-14-yn-5-one (15)** To a 0 °C solution of **13** (4.8 g, 13.4 mmol, 1 equivalent) in DCM (120 mL), 2,3-dichloro-5,6-dicyano-1,4-benzoquinone (6.082 g, 26.8 mol, 2 equivalents) was added as a solid, followed by water (15 mL). The reaction was stirred for 30 min at 0 °C and then for 1 h at room temperature. Once the reaction had gone to completion, by TLC (3:1 hexanes : ethyl acetate, Rf = 0.2, KMnO_4_ staining) The reaction was then poured over 500 mL saturated sodium bicarbonate, extracted 2 x 200 mL DCM, then washed 2 x 300 mL with 10% sodium bicarbonate and 2 x 300 mL brine. The crude orange residue was purified by flash chromatography (gradient of 30 % ethyl acetate to 50 % ethyl acetate in hexanes) yielding **15** (2.6 g, 17.17 mmol) as a pale orange semi-crystalline solid in 59 % yield. ^1^H NMR (400 MHz, CDCl_3_) δ = 3.62 (t, J = 6.4, 2H), 2.47 (t, J = 7.2, 2H), 2.40 (t, J = 7.2, 2H), 2.17 (td, J = 7.2, J = 2.8, 2H), 1.94 (t, J = 2.8, 1H), 1.68-1.64 (m, 2H), 1.57-1.41 (m, 6H), 1.42-1.37 (m, 2H), 1.28-1.15 (m, 6H); ^13^C NMR (400 MHz, CDCl_3_) δ = 2.11.6, 84.8, 68.1, 62.3, 42.8, 42.3, 32.2, 29.3, 28.9, 28.7, 28.4, 23.9, 19.7, 18.4

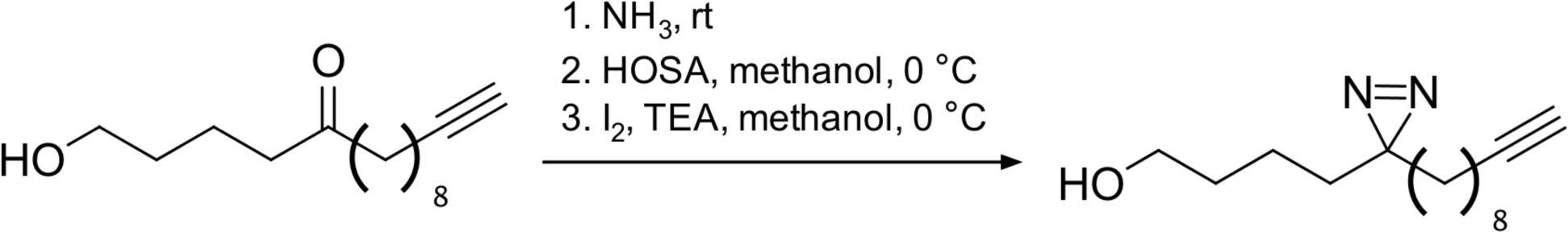

**4-[3-(Dec-9-yn-1-yl)-3H-diazirin-3-yl]butan-1-ol (17) 26** (440 mg, 1 equivalent) was placed in an oven-dried bomb flask and cooled to -85 °C. Ammonia (25 mL) was condensed on top of the compound and the bomb flask was capped. The mixture was allowed to warm to room temperature, stirring, whereupon 17 dissolved in the ammonia. The reaction stirred for 24hr at room temperature, and then the ammonia was evaporated overnight, into an aqueous 10% citric acid bath. Hydroxylaminesulfonic acid (459 mg, 2.2 equivalents) was dissolved in 15 mL anhydrous methanol and added to the residue in the bomb flask at 0 °C. This was stirred for 1 hr at rt, and then the reaction mixture was filtered to remove white solids and the methanol was evaporated. The residue was dissolved in 50 mL ethyl acetate and washed 1x 25 mL 10% citric acid and 1x25 mL brine, dried over MgSO4, and concentrated. The crude residue was oxidized without further purification, by dissolving in 50 mL methanol, cooling to 0 °C, adding TEA (4.1 mL, 2 equivalents), and I_2_ (468 g, 1 equivalent) portion wise as a solid until the brown iodine color persisted. Methanol was once again evaporated, the residue was dissolved in 50 mL ethyl acetate and washed 4x 50mL with sodium thiosulfate and 1x 50mL brine. TLC (2:1 hexanes: ethyl acetate, potassium permanganate staining) indicated a complex mixture of products; isolation of the spot with Rf = 0.45 resulted in the correct product. The crude reside was purified by flash chromatography (9:1 hexanes : ethyl acetate) yielding **17** (97.5 mg) in 21 % yield as a pale yellow solid. ^1^H NMR (400 MHz, CDCl_3_) δ = 3.54 (t, J = 6.4, 2H), 2.11 (td, J = 7.2, J= 2.8, 2H), 1.87 (t, J = 2.8, 1H), 1.68-1.64 (m, 2H), 1.33-1.27 (m, 6H), 1.21-1.15 (m, 6H), 1.14-1.08 (m, 2H), 1.03-0.97 (m, 2H); ^13^C NMR (400 MHz, CDCl_3_) δ = 84.8, 77.2, 68.1, 62.6, 32.8, 32.7, 32.2, 29.2, 29.1, 28.9, 28.7, 28.4, 23.8, 20.2, 18.4

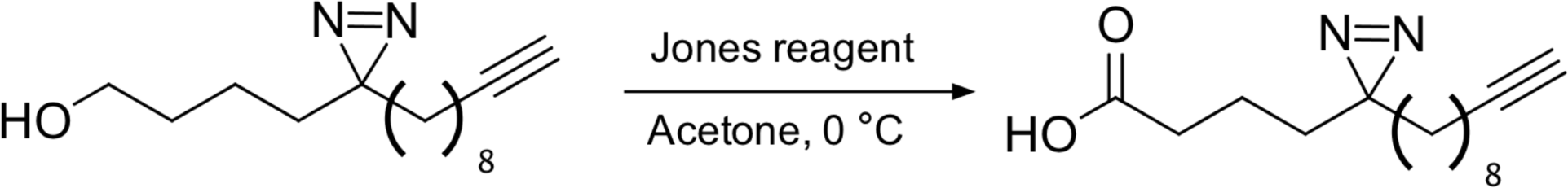

**4-[3-(Dec-9-yn-1-yl)-3H-diazirin-3-yl] butanoic acid (19) 17** (94 mg, 1 equivalent) was dissolved in acetone (20 mL) and cooled to 0 °C. Jones reagent (2 M, 200µL, 1 equivalent) was added, dropwise, until the pink color of the reagent persisted. The reaction was stirred for 15 min at 0°C, and after the appearance of product by TLC (2:1 hexanes : ethyl acetate, Rf = 0.3, KMnO_4_ staining) then quenched with isopropanol, producing a bright, blue-green precipitate. The precipitate was filtered and then the acetone was removed under reduced pressure. The residue was dissolved in ethyl acetate (20 mL) and washed 2x with saturated sodium bicarbonate and 1x brine, dried over magnesium sulfate, and concentrated, yielding pure **19** (55.9 mg, 56 %) as a white solid. ^1^H NMR (400 MHz, CDCl_3_) δ = 2.25 (t, J = 7.2, 2H), 2.11 (td, J = 7.2, J = 2.8, 2H), 1.87 (t, 2.8, 1H), 1.46-1.14 (m, 16H), 1.14-0.99 (m, 2H); ^13^C NMR (400 MHz, CDCl_3_) δ = 178.3, 84.8, 68.1, 33.1, 32.7, 32.2, 29.7, 29.2, 29.1, 28.9, 28.7, 28.4, 23.8, 19.0, 18.4

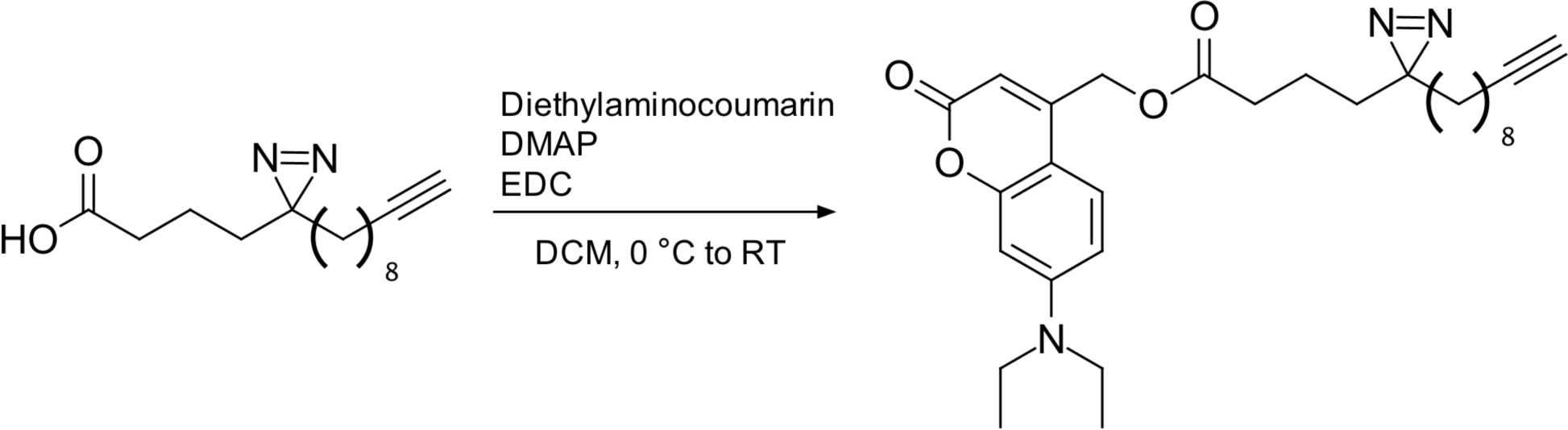

**[7-(Diethylamino)-2-oxo-2H-chromen-4-yl] methyl 4-[3-(dec-9-yn-1-yl)-3H-diazirin-yl] butanoate (3-8 Fatty Acid, 3) 19 (**54 mg, 1 equivalent) was dissolved in 10 mL DCM. EDC (78.3 mg, 2 equivalents) and DMAP (4.99 mg, 0.2 equivalents) were added and stirred together for 15 min, as the mixture gradually turned orange. 7-(Diethylamino) 4-hydroxymethylcoumarin (55.6 mg, 1.1 equivalents) was dissolved in 5 mL DCM. The coumarin and the activated compound **19** were both cooled to 0 °C and the coumarin added was added to compound **19**. The reaction was allowed to warm to room temperature overnight. Appearance of product was observed by TLC (3:1 hexanes : ethyl acetate, Rf = 0.1, potassium permanganate and UV visualization), and the reaction was poured over citric acid, extracted 2 x 15 mL with DCM, washed once with brine, and concentrated under reduced pressure. The crude residue was purified initially by flash chromatography (gradient of 5 % to 10 % ethyl acetate in hexanes) to produce 7 as a pale yellow solid (130 mg, 64.9 % yield). ^1^H NMR (400 MHz, CDCl_3_) δ = 7.21 (d, J = 8.8, 1H), 6.51 (dd, J = 8.8, J = 2.8, 1H), 6.45 (d, J = 2.8, 1H), 6.04 (s, 1H), 5.15 (s, 2H), 3.35 (q, J = 7.2, 4H), 2.34 (t, J = 7.2, 2H), 2.1 (td, J = 7.2, J = 2.8, 2H), 1.87 (t, J = 2.8, 1H), 1.48-1.14 (m, 4H), 1.37-1.16 (m, 12H), 1.14 (t, J = 7.2, 6H), 1.03-0.98 (m, 2H); ^13^C NMR (400 MHz, CDCl_3_) δ = 172.3, 161.9, 156.3, 150.7, 149.3, 124.4, 108.7, 106.5, 106.0, 97.9, 84.7, 68.1, 61.4, 44.8, 33.3, 32.7, 32.3, 29.2, 29.1, 28.9, 28.7, 28.4, 28.2, 23.8, 19.2, 18.4, 12.4

#### Cell culture

Huh7 cells were maintained in standard tissue-culture treated vessels in DMEM supplemented with 10% FBS, 1% non-essential amino acids and 1% penicillin-streptomycin at 37 °C and 5% CO_2_.

#### Antibodies and chemicals

- Antibodies

- anti-Giantin (mouse, abcam ab37266)
- anti-PDI (rabbit, Cell Signaling 3501S)
- Goat anti-rabbit IgG (H+L) Alexa Fluor Plus 488 (Invitrogen A32731TR)
- Goat anti-Mouse IgG (H+L) Alexa Fluor 555 (Invitrogen A-21422)
- anti-PITPNB (rabbit, Proteintech, 13110)
- Anti-rabbit IgG, HRP-linked Antibody (Cell Signaling, #7074)
- Chemicals

- A647 picolyl azide (Vector laboratories CCT-1300)
- Picolyl azide-agarose beads (Vector laboratories CCT-1408)
- Biotin azide (Vectory laboratories CCT-1265)
- Monomeric avidin agarose resin (Thermo Scientific 20228)

#### Uncaging and photocrosslinking

For uncaging and photocrosslinking, two lamps were primarily used: a UV type 1 LED lamp (Nailstar model NS-02, https://www.amazon.de/gp/ product/B01286DTFQ/ref=ppx_yo_dt_b_asin_title_o02_s00?ie=UTF8&psc=1) for uncaging — “Uncaging Lamp”, and a 36W/365 nm lamp (MelodySusie, Pro04 model, https://www.amazon.com/MelodySusie-Professional-Setting-Manicure-Pedicure/dp/B012MEZP2E?ref_=ast_sto_dp&th=1&psc=1) for photocrosslinking — “Photocrosslinking Lamp”. Unless stated otherwise below, each lamp was used for 5 minutes. For the lamp comparison experiment (see below), an additional lamp was used, a 1,000 W high-power Mercury-Xenon lamp (Newport), fitted with either a 400 nm high-pass filter (for uncaging) or a 345 nm high-pass filter (for photocrossinking).

#### Lamp comparison

Huh7 cells were seeded in 24-well plates containing glass coverslips and grown to 75% confluence. 8-3 FA was diluted to 50 µM in complete media, 200 µL probe solution was added to each well, and was allowed to sit on cells for 30 min at 37 °C prior to uncaging. Cells were uncaged either with the Uncaging Lamp or with the Newport lamp fitted with a 400 nm high-pass filter, for either 2 min or 5 min, and then photocrosslinked either with the Photocrosslinking Lamp or with the Newport lam fitted with a 345 nm high-pass filter. A “no light” background control was also included, which had been treated with 8-3 FA in the same manner as above but kept in the dark prior to fixation. Cells were fixed in methanol for 20 min. Cells were washed three times with PBS to remove organic solvent, and then 200 µL (for 24-well plates) of a click mix were added (1 mM copper sulfate, 1 mM sodium ascorbate, 100 µM TBTA, 2 µM A647 picolyl azide, in PBS). The reaction was allowed to proceed for 1 h in the dark. Cells were stained with DAPI (1:1000 in PBS) for 10 minutes. Coverslips were mounted on slides and imaged on a Zeiss LSM 980 Laser-Scanning 4-channel confocal microscope.

#### Analysis of trifunctional lipids by thin-layer chromatography

*Probe labeling* Huh7 cells were seeded in 6-well plates and grown to 75% confluence. Dilutions of trifunctional probe were made in complete media, for a final concentration of 50 µM for each fatty acid, with a corresponding DMSO control. 1 mL probe dilution was added to each plate and allowed to sit on cells for 30 min at 37 °C prior to uncaging. Dishes were exposed to 400 nm light via the Uncaging Lamp for five min to uncage the probe. Probe solutions were removed and replaced with DMEM, and then cells were returned to the incubator for 1 or 24 h to allow for metabolism. *Lipid extraction* Lipids were extracted with a modified Bligh-Dyer extraction; briefly, dishes were rinsed 4x with PBS, and 1 mL of a 2 : 0.8 methanol:water mixture was added, and cells were scraped into this mixture and moved to a glass tube, to which 1 mL of chloroform was added. Tubes were vortexed to mix, and the layers were allowed to separate at -20 °C for 1 h to overnight. Tubes were centrifuged (1,000 x g for 10 min) to ensure complete separation of layers, and then the chloroform layer was removed to a fresh tube and then dried under a stream of nitrogen. *Click labeling* Extracted lipids were re-dissolved in 10 µL chloroform, and 40 µL of a click mix (5 µL each of 1 mM TBTA, 10 mM copper sulfate, 10 mM sodium ascorbate, and 10 mM 3-azido-7-diethylaminocoumarin, and 20 µL ethanol) was added. The reaction was allowed to proceed for 3 h in the dark, and then extracts were once more dried under a stream of nitrogen. *Plating and running TLC* Extracted lipids were re-dissolved in 10 µL chloroform and plated on 10x10 cm HPTLC silica 60 Å glass plates without F254 fluorophore. Lipids were resolved by a 2-step system: first using chloroform/methanol/saturated aqueous ammonium hydroxide 65:25:4 for 6cm, then drying, and then using hexanes/ethyl acetate 1:1 for 9cm. Fluorescently labeled lipids were visualized using a Sapphire molecular imager in the Cy2 channel. Images were processed in Fiji software to subtract background via the rolling-ball method with radius = 40 px.

#### Subcellular visualization of lipids by confocal microscopy

*Probe labeling* Huh7 cells were seeded on coverslips in 24-well plates, and grown to about 70% confluence. Dilutions of trifunctional probe were made in complete media to a final concentration of 50 µM for each fatty acid. 200 µL of probe dilution was added to each plate and allowed to sit on cells for 30 min at 37 °C prior to uncaging. Dishes were exposed to 400 nm light for fives minutes to uncage the probe, using the Uncaging Lamp and returned to the incubator for 5, 30, or 60 min to allow for metabolism, then exposed to 350 nm light for five minutes to photo-crosslink the probe, using the Photocrosslinking Lamp and immediately fixed. Cells were fixed by washing twice with PBS, then left in methanol for 20 min. *Click labeling* Cells were washed three times with PBS to remove organic solvent, and then 200 µL (for 24-well plates) or 70 µL (for 96-well plates) of a click mix were added (1 mM copper sulfate, 1 mM sodium ascorbate, 100 µM TBTA, 2 µM A647 picolyl azide, in PBS). The reaction was allowed to proceed for 1 h in the dark. *Antibody staining* Click mix was removed, cells were washed twice with PBS, and blocking buffer (2% BSA, 0.1% Triton-X-100 in PBS) was added. Cells were blocked for 1 h before the addition of primary antibodies. Primary antibodies (anti-Giantin, anti-PDI, catalogue numbers listed above) were diluted 1:250 in blocking buffer and left on cells, with rocking, overnight at 4 °C. The next day, primary antibodies were removed, cells were washed three times with PBS, and fluorescent secondary antibodies were added, either A488 anti-rabbit or A555 anti-mouse, 1:500 dilution in blocking buffer, for 1 h at room temperature, with rocking. Secondary antibodies were removed, cells were washed three times with PBS, and DAPI (1:1000) was added for ten min. Cells were imaged within a week after staining, on a Zeiss LSM 980 Laser-Scanning 4-channel confocal microscope with Airyscan.2*^1^*. *Image analysis* Pearson’s correlation coefficients between the lipid signal and the signal for each organelle marker were calculated using a CellProfiler pipeline^2^. Individual cells were selected based off of regions of intensity of the lipid signal and coefficients were calculated within each cell.

#### Isolation and identification of protein-lipid complexes by LC-MS/MS

*Probe labeling* Huh7 cells were seeded in 10cm dishes and grown to 90% confluence. Dilutions of trifunctional probe were made in complete media to a final concentration of 50 µM for each fatty acid. 3 mL of probe dilution was added to each plate and allowed to sit on cells for 30 min at 37 °C prior to uncaging for 5 minutes with the Uncaging Lamp. Cells treated with fatty acids were subjected to photocrosslinking for five minutes with the Photocrosslinking Lamp 1 h after uncaging. After photocrosslinking, cells were washed three times with PBS and scraped into 2 mL of ice-cold PBS. Cells were pelleted by centrifugation (1,000 x g for 5 min), supernatant was decanted, and cells were resuspended in 500 µL PBS. *Sample preparation for proteomics* Cells were lysed by probe sonication, on ice, in three 15-second bursts. Lysates were subjected to a click reaction with picolyl azide agarose beads: 200 µL azide beads were washed once in DI water, then added to the cell lysate, along with copper sulfate (1mM, final concentration), sodium ascorbate (1 mM, final concentration), and TBTA (100 µM, final concentration). Samples were rotated at room temperature for 1 h. Beads were spun down (1,000 x g for 2 min), transferred to 2 mL centrifuge columns, and washed extensively: 3x with PBS, 5x with bead wash buffer 1 (100 mM Tris-HCl, pH = 8.0, 250 mM NaCl, 5 mM EDTA, 1% SDS), 10x with bead wash buffer 2 (100 mM Tris-HCl, pH = 8.0, 8M urea). Beads were transferred from the column in PBS to a clean eppendorf tube and spun down. Isolated proteins were reduced (by resuspending them in 1 mM digestion buffer [100 mM Tris-HCl, pH = 8.0, 2 mM CaCl_2_, 10% ACN], adding DTT to 10 mM, and incubating at 42 °C for 30 min), alkylated (by spinning down beads and resuspending them in 1 mL 40 mM aqueous iodoacetamide and incubating them at room temperature in the dark for 30 min). Bead-bound proteins were then digested by spinning them down, adding 50 µL of digestion buffer and 1 µL of LC-MS grade trypsin, and shaken at 37 °C overnight. Peptides were then desalted on C18 columns and the eluent was frozen at -80 °C. *Identification of isolated proteins by LC-MS/MS.* Dried peptides were shipped to the EMBL proteomics core facility where they were TMT-labeled using the TMT-16-plex system and analyzed by LC-MS/MS on an Orbitrap Fusion Lumos mass spectrometer (Thermo Scientific). Peptides were separated using an Ultimate 3000 nano RSLC system (Dionex) equipped with a precolumn (C18 PepMam100, 5mm, 300 µm i.d., 5 µm, 100 Å) and an analytical column (Acclaim PepMap 100. 75 x 50 cm C18, 3 mm, 100 Å) connected to a nanospray-Flex ion source. The peptides were loaded onto the precolumn in solvent A (0.1% formic acid) and eluted in a gradient of solvent B (0.1% formic acid in acetonitrile) from 2 to 85% over 120 min at 0.3 µL per minute. Data were collected in positive ion mode with a spray voltage of 2.4 kV and capillary temperature of 275 °C. Full scan MS spectra with a mass range of 300–1500 m/z were acquired in profile mode using a resolution of 120,000. AGC target was set to 50% and a max injection time of 250 ms. Precursors were isolated using the quadrupole with a window of 1.4 m/z and fragmentation was triggered by HCD in fixed collision energy mode with fixed collision energy of 30%. MS2 spectra were acquired with the Orbitrap with a resolution of 15.000. Normalized AGC target was set to 200% and a max injection time of 32 ms. *Analysis of proteomics data.* Acquired data were analyzed using IsobarQuant^3^ and Mascot V.24 (Matrix Science) using a reverse UniProt FASTA database (UP000005640) including common contaminants. Only proteins that were quantified with two unique peptide matches in both replicates were kept for the analysis. A variance stabilization normalization was performed on the log2 raw data^4^, and enrichments of proteins in the (+) UV condition over the (-) UV condition was calculated using LIMMA analysis^5^. A protein is considered a “hit” in a certain condition if the false discovery rate is smaller than 0.05 and the fold change is at least 2; a protein is considered a “candidate” if a the false discovery rate is smaller than 0.2 and the fold change is at least 1.5.

#### In gel fluorescence of protein-lipid complexes

*Probe labeling* Huh7 cells were seeded in 10cm dishes and grown to 90% confluence, in triplicate for each condition. Dilutions of trifunctional probe were made in complete media to a final concentration of 50 µM for each fatty acid. 3 mL of probe dilution was added to each plate and allowed to sit on cells for 30 min at 37 °C prior to uncaging for 5 minutes with the Uncaging Lamp. Cells treated with fatty acids were subjected to photocrosslinking for five minutes with the Photocrosslinking Lamp 1 hour after uncaging. (-) UV controls for each probe were prepared by keeping probe-treated cells in the dark prior to harvest. After photocrosslinking, cells were washed three times with PBS and scraped into 2 mL of ice-cold PBS. Cells were pelleted by centrifugation (1,000 x g for 5 min), supernatant was decanted, and cells were resuspended in 250 µL PBS. *Lysis, click reaction, and gel running* Cells were lysed by probe sonication, on ice, in three 15-second bursts. Lysates were subjected to a BCA assay to determine protein concentration, and amount of protein in each condition were normalized. 25 µL of lysate, diluted in PBS to produce a uniform 32.5 µg of protein per condition, was subjected to a click reaction with A647 picolyl azide (by adding 2 µL of a master mix containing the azide, copper sulfate, sodium ascorbate, and TBTA, for a final concentration of 20 µM azide, 100 µM TBTA, 1 mM copper sulfate, and 1 mM sodium ascorbate), for 1 hour at room temperature, in the dark. Proteins were gently solubilized for the gel by incubating with Laemmli buffer for 30 minutes at 60 °C, and then run on a 12.5% SDS-PAGE gel. The far-red fluorescence was visualized on an Azure Sapphire Biomolecular Imager. Following fluorescent visualization, the gel was stained with Blazin’ Blue Protein Gel Stain to verify the uniformity of protein amount between samples.

#### Biotin pulldown and western blotting of protein-lipid complexes

*Pulldown* The same lysates prepared in the “in gel fluorescence” section above were used for pulldown experiments. Protein amounts were normalized by dilution with PBS based on the BCA assay, with a total of 260 µg of protein per sample, in 500 µL total. Lysates were subjected to a click reaction with biotin azide (by adding 5.15 µL of a master mix containing the azide, copper sulfate, sodium ascorbate, and TBTA, for a final concentration of 5 µM azide, 100 µM TBTA, 1 mM copper sulfate, and 1 mM sodium ascorbate), for 2.5 hours at room temperature, rocking. Excess biotin azide was removed on three sequential Zeba desalting columns (7K MWCO): columns were equilibrated three times with PBS, and the click reaction was loaded onto the column and spun through following manufacturer’s instructions. Flow-through from the first column was loaded onto the second column, and so on. Flow-through from the third column was incubated overnight at 4 °C with 200 µL monomeric avidin agarose resin. In the morning, beads were washed four times with 1 mL of a wash buffer (0.05% Triton-X-100 in PBS), spinning at 2,500 x g for 2 minutes after each wash. Proteins were eluted off beads by adding 60 µL of 1x Laemmli buffer and incubating at 60 °C for 30 minutes. *Western blotting* Eluent from the agarose beads, as well as the input whole-cell lysate, was run on a 12.5% SDS-PAGE gel and then transferred to a PVDF membrane. After blocking in 3% BSA in TBST, the membrane was treated with anti-PITPNB primary antibody (1:1000) overnight at 4 °C, and then an anti-rabbit HRP secondary antibody (1:5000) for 1 hour at room temperature, before treatment with a chemiluminescent substrate and visualization on an ImageQuant 4000 imager.

## SUPPLEMENTAL FIGURES

**Supplementary Figure 1:**
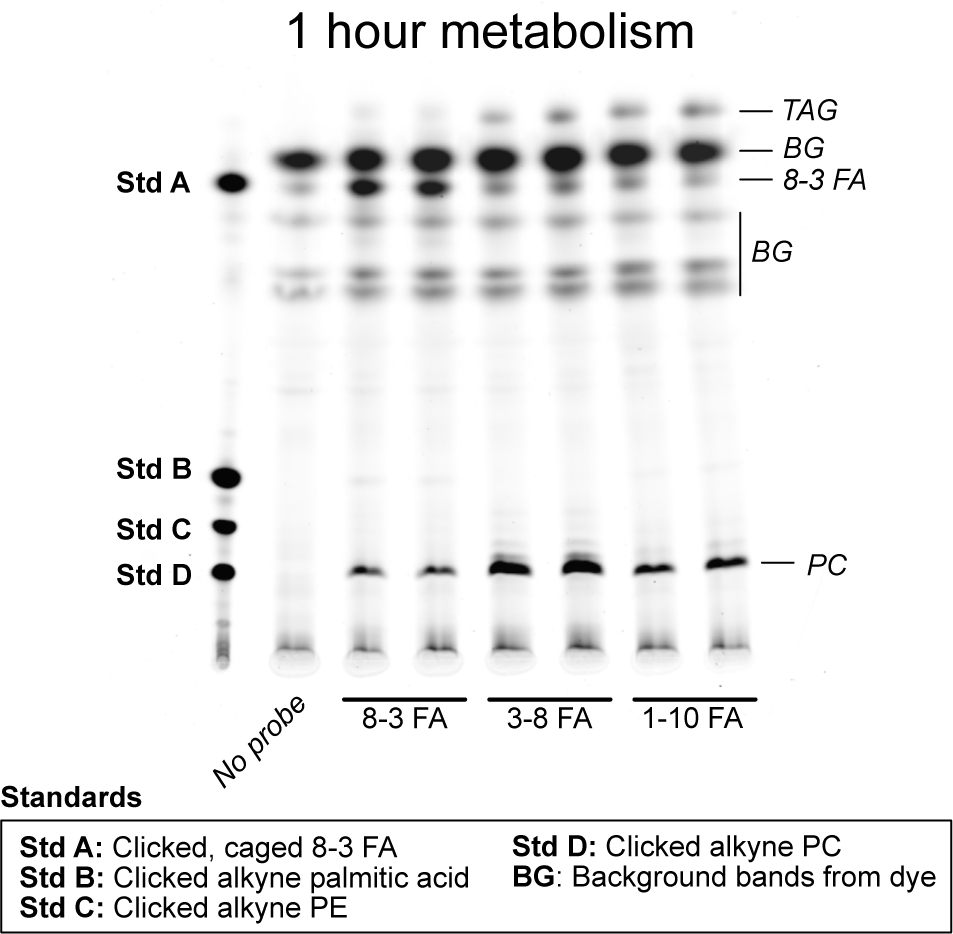
Metabolism of trifunctional fatty acids TLC of lipid extracts from Huh7 cells treated with 8-3 FA, 3-8 and 1-10 FA and harvested 1 hour after uncaging. TAG = triacylglyerol; BG = background; PE = phosphatidylethanolamine; PC = phosphatidylcholine.

**Supplementary Figure 2:**
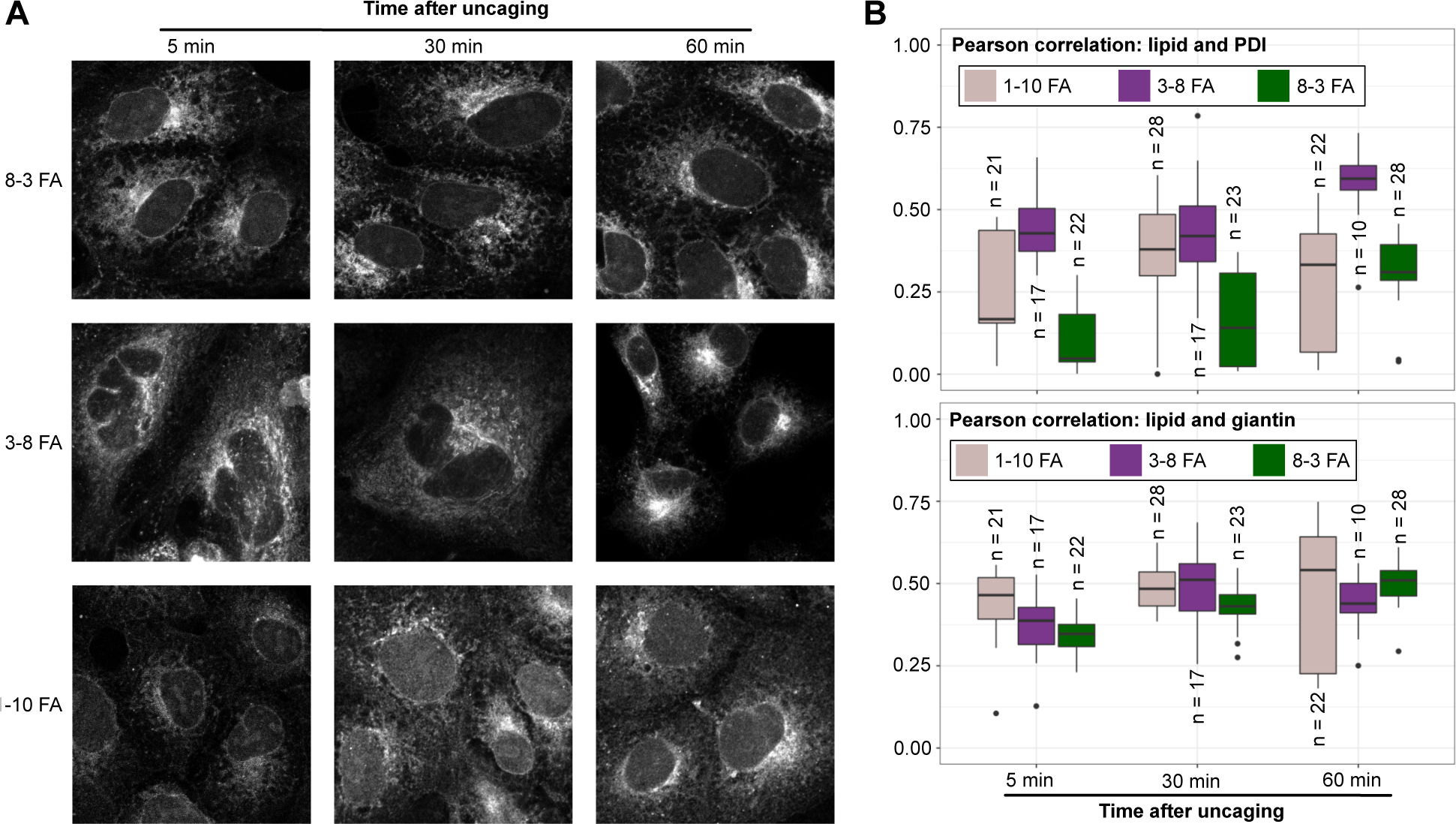
Subcellular localization of trifunctional fatty acids. (**A**) Representative images of Huh7 cells treated with 8-3 FA, 3-8 FA, or 1-10 FA, exposed to 400nm light to uncage the probe, and crosslinked with 350nm light 5, 30, or 60 minutes after uncaging, and subjected to click reactions with A647 picolyl azide. Representative images of the far red signal at each timepoint are shown; images are representative of four biological replicates over two independent experiments (**B**) Colocalization of each fatty acid with markers for the ER (PDI) or Golgi (Giantin). Pearson coefficients were calculated for individual cells using a Cellprofiler pipeline; the number of cells used for each condition are indicated by their respective boxplots.

**Supplementary Figure 3.**
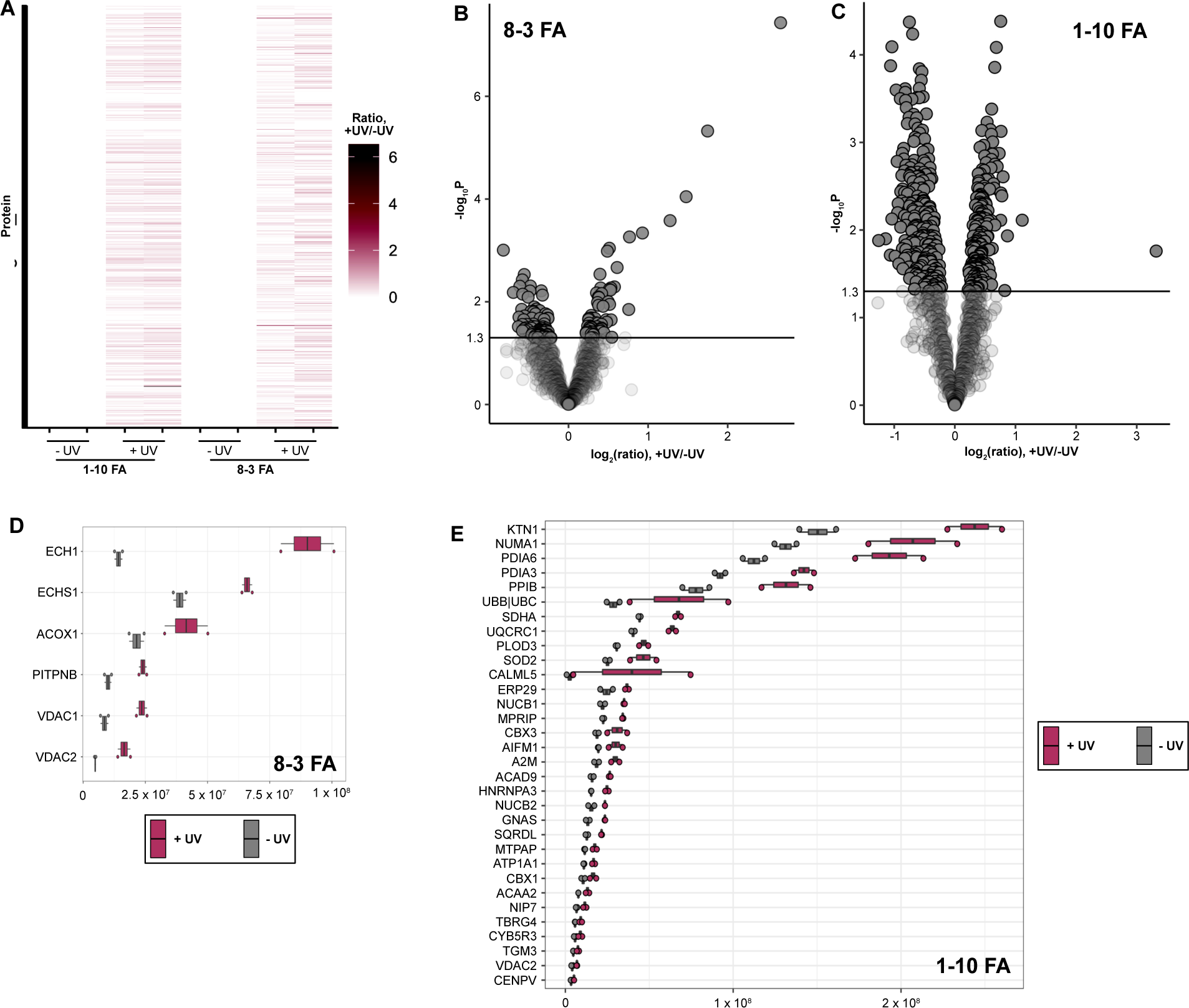
Overview of proteomic analysis. Huh7 cells were treated with 8-3 FA or 1-10 FA, uncaged, and photocrosslink 1 hour after uncaging (+ UV), or uncaged without subsequent photocrosslinking (-UV). Cell extracts were lysed by probe sonication and subjected to a click reaction with azide agarose prior to washing to removing nonbound proteins and trypsin digestion. Tryptic digests were desalted and analyzed by LC-MS/MS. (**A**) Heat map of the log-transformed ratio of the intensity of each protein in the +UV sample versus the intensity for that protein in the - UV sample (**B**) Volcano plot of negative log-transformed p-values and fold changes (+UV over -UV) from the Limma analysis of 8-3 FA (**C**) Volcano plot of negative log-transformed p-values and fold changes (+UV over -UV) from the Limma analysis of 1-10 FA (**D**) Normalized intensities for the “hit” and “candidate” proteins in the 8-3 FA-treated samples (**E**) Normalized intensities for the “hit” and “candidate” proteins in the 1-10 FA-treated samples

**Supplementary Figure 4.**
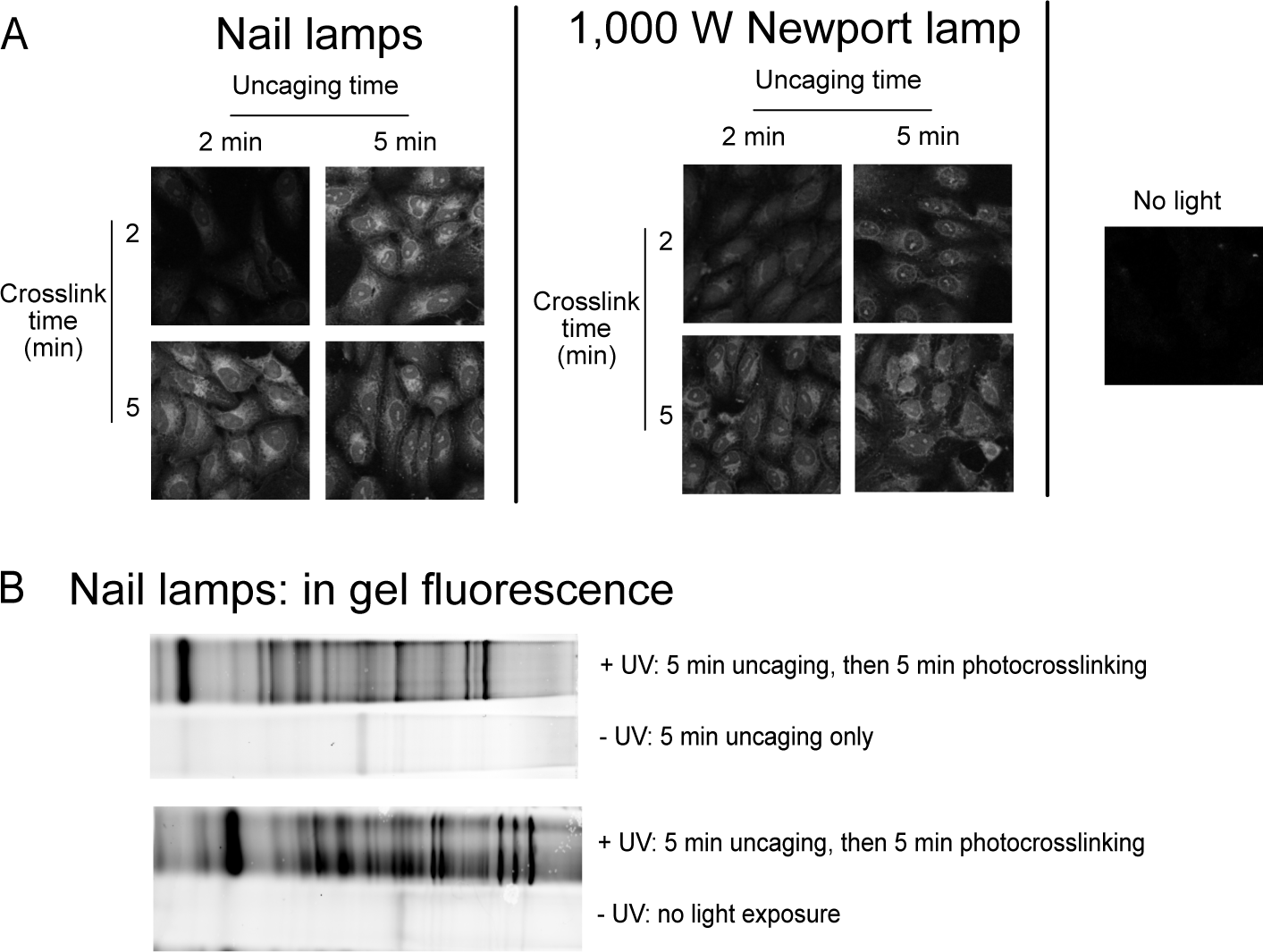
Lamp comparison. (**A**) Confocal microscopy of cells treated with 8-3 FA and then uncaged and photocrosslinked for the indicated times with either the nail lamps described in the Supplementary Methods or a 1,000W Mercury-Xenon light source fitted with 400 nm (for uncaging) or 345 nm (for photocrosslinking) high-pass filters. (**B**) In-gel fluorescence of lysates from cells treated with 8-3 FA and then uncaged and photocrosslinked for 5 minutes with the nail lamps; negative controls are either 1) uncaging for five minutes only (no photocrosslinking) or 2) no light exposure, maintained in the dark until cells were scraped for lysis

**Supplementary Figure 5.**
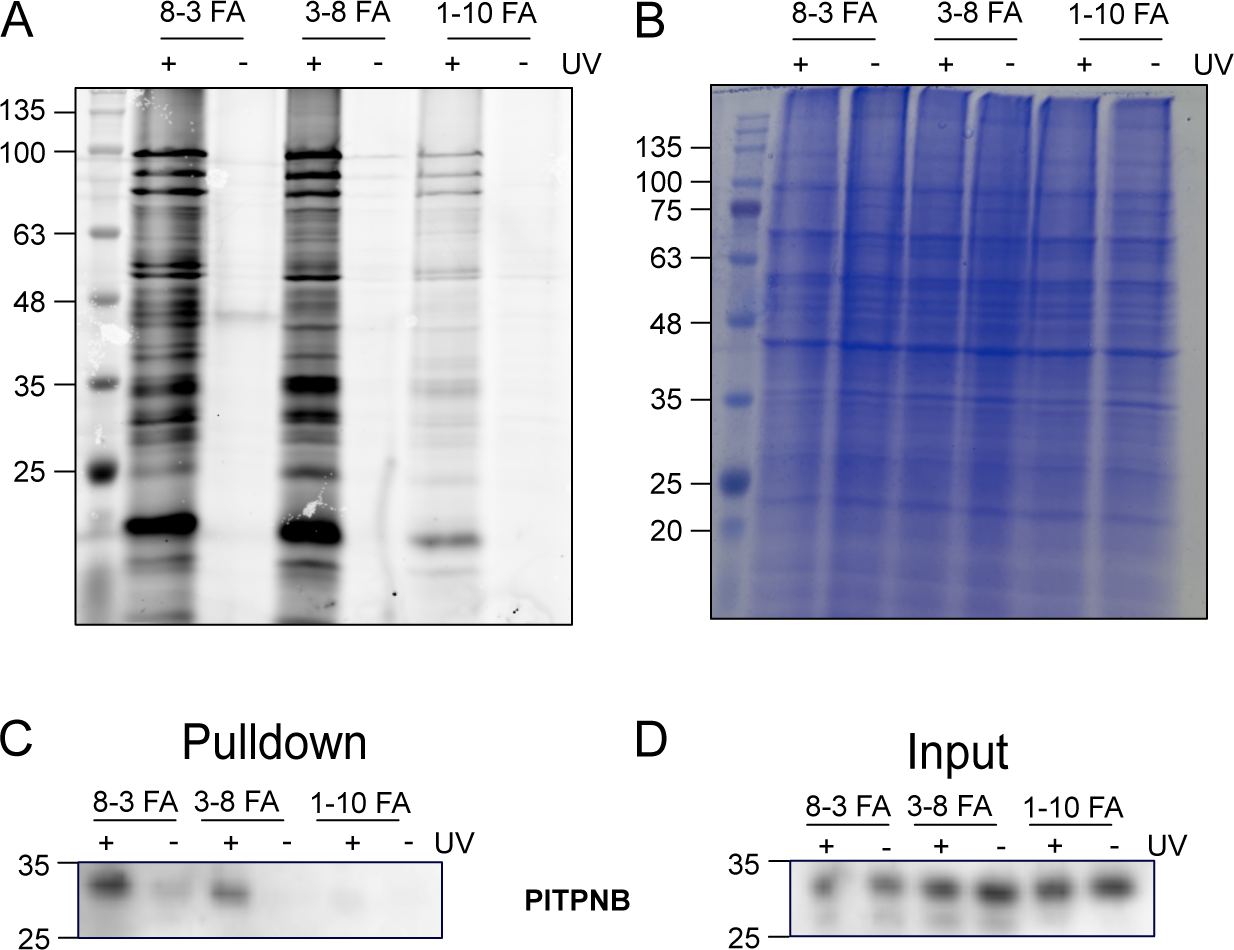
In-gel fluorescence and biotin pulldowns. (**A**) In-gel fluorescence of lysates from Huh7 cells treated with the indicated probe, in the presence or absence of UV irradiation (from “Photocrosslinking Lamp”). Shown is fluorescence from an A647 picolyl azide clicked to the lysate. (**B**) Blazin’ Blue staining of total protein from the gel in A. (**C**) PITPNB staining of protein-lipid complexes subjected to a click reaction with biotin azide, pulled down with monomeric avidin agarose, and eluted with Laemmli buffer. Image is representative of three independent experiments. (**D**) PITPNB staining of the input lysate that was then subjected to the pulldown in (C)

## REFERENCES

1. Watschinger, K.; Keller, M. A.; McNeill, E.; Werner, E. R., Proc Natl Acad Sci U S A, 2015, 112 (8), 2431–2436.

2. Maekawa, M.; Fairn, G. D., J. Cell. Sci. 2014, 127, 4801–4812.

3. Thiele, C.; Papan, C.; Hoelper, D.; Kusserow, K.; Gaebler, A.; Schoene, M.; Piotrowitz, K.; Lohmann, D.; Spandl, J.; Stevanovic, A.; Shevchenko, A.; Kuerschner, L. ACS Chem Biol 2012, 7 (12), 2004–11.

4. Gaebler, A.; Milan, R.; Straub, L.; Hoelper, D.; Kuerschner, L.; Thiele, C., J Lipid Res 2013, 54 (8), 2282–2290.

5. Haberkant, P.; Holthuis, J. C. M., Biochim Biophys Acta 2014, 1841 (8), 1022–1030.

6. Farley, S.; Laguerre, A.; Schultz, C., Curr Opin Chem Biol 2021, 65, 42–48.

7. Hoglinger, D.; Haberkant, P.; Aguilera-Romero, A.; Riezman, H.; Porter, F. D.; Platt, F. M.; Galione, A.; Schultz, C., Elife 2015, 4, e10616.

8. Hoglinger, D.; Nadler, A.; Haberkant, P.; Kirkpatrick, J.; Schifferer, M.; Stein, F.; Hauke, S.; Porter, F. D.; Schultz, C., Proc Natl Acad Sci U S A 2017, 114 (7), 1566–1571.

9. Muller, R.; Citir, M.; Hauke, S.; Schultz, C., Chem. Eur. J. 2020, 26 (2), 384–389.

10. Muller, R.; Kojic, A.; Citir, M.; Schultz, C., Angew Chem Int Ed Engl 2021, 60 (36), 19759–19765.

11. Farley, S.; Stein, F.; Haberkant, P.; Tafesse, F. G.; Schultz, C., ACS Chem Biol 2024, 19 (2), 336–347

12. Haberkant, P.; Raijmakers, R.; Wildwater, M.; Sachsenheimer, T.; Brugger, B.; Maeda, K.; Houweling, M.; Gavin, A. C.; Schultz, C.; van Meer, G.; Heck, A. J.; Holthuis, J. C., Angew Chem Int Ed Engl 2013, 52 (14), 4033–8.

13. Brown, C. A.; Yamashita, A., JACS, 1975, 97 (4), 891–892.

14. Avocetien, K.; Li, Y.; O’Doherty, G. A.; Trost, B. M.; Li, C.-J., Eds. Wiley-VCH Verlag GmbH & Co. KGaA:2014; pp 365-394.

15. Carrasco, S.; Merida, I., Trends Biochem Sci 2007, 32 (1), 27–36.

16. Dunn, K. W.; Kamocka, M. M.; McDonald, J. H., Am J Physiol Cell Physiol 2011, 300 (4), C723–42.

17. McQuin, C.; Goodman, A.; Chernyshev, V.; Kamentsky, L.; Cimini, B. A.; Karhohs, K. W.; Doan, M.; Ding, L.; Rafelski, S. M.; Thirstrup, D.; Wiegraebe, W.; Singh, S.; Becker, T.; Caicedo, J. C.; Carpenter, A. E., PLoS Biol 2018, 16 (7), e2005970.

18. Huber, W.; von Heydebreck, A.; Sültmann, H.; Poustka, A.; Vingron, M., Bioinformatics 2002, 18 Suppl 1:S96–104.

19. Franken, H.; Mathieson, T.; Childs, D.; Sweetman, G. M. A.; Werner, T.; Tögel, I.; Doce, C.; Gade, S.; Bantscheff, M.; Drewes, G.; Reinhard, F. B. M.; Huber, W.; Savitski, M. M., Nat Protocols 2015, 10 (10), 1567–1593.

20. Fedoryshchak, R. O.; Goerlik, A.; Shen, M.; Shchepinova, M. M.; Pérez-Dorado, I.; Tate, E. W., Chem. Sci 2023, 14, 2419–2430.

## REFERENCES - METHODS

1. Huff, J., The Airyscan detector from ZEISS: confocal imaging with improved signal-to-noise ratio and super-resolution. Nature Methods 2015, 12 (12), i-ii.

2. McQuin, C.; Goodman, A.; Chernyshev, V.; Kamentsky, L.; Cimini, B. A.; Karhohs, K. W.; Doan, M.; Ding, L.; Rafelski, S. M.; Thirstrup, D.; Wiegraebe, W.; Singh, S.; Becker, T.; Caicedo, J. C.; Carpenter, A. E., CellProfiler 3.0: Next-generation image processing for biology. PLoS Biol 2018, 16 (7).

3. Franken, H.; Mathieson, T.; Childs, D.; Sweetman, G. M. A.; Werner, T.; Tögel, I.; Doce, C.; Gade, S.; Bantscheff, M.; Drewes, G.; Reinhard, F. B. M.; Huber, W.; Savitski, M. M., Thermal proteome profiling for unbiased identification of direct and indirect drug targets using multiplexed quantitative mass spectrometry. Nature Protocols 2015, 10 (10), 1567–1593.

4. Huber, W.; von Heydebreck, A.; Sueltmann, H.; Poustka, A.; Vingron, M., Variance Stabilization Applied to Microarray Data Calibration and to the Quantiication of Differential Expresion. Bioinformatics 2002, 18 Suppl. 1, S96–S104.

5. Ritchie, M. E.; Phipson, B.; Wu, D.; Hu, Y.; Law, C. W.; Shi, W.; Smyth, G. K., limma powers differential expression analysis for RNA-sequencing and microarray studies. Nucleic Acids Res 2015, 43 (7).

